# Telomerase RNA component knockout exacerbates *S. aureus* pneumonia by extensive inflammation and dysfunction of T cells

**DOI:** 10.1101/2024.06.19.599722

**Authors:** Yasmina Reisser, Franziska Hornung, Antje Häder, Thurid Lauf, Sandor Nietzsche, Bettina Löffler, Stefanie Deinhardt-Emmer

**Affiliations:** Institute of Medical Microbiology, Jena University Hospital, Am Klinikum 1, 07747 Jena, Germany; Else Kröner Graduate School for Medical Students “JSAM”, Jena University Hospital, 07747 Jena, Germany; Center for Electron Microscopy, Jena University Hospital, Ziegelmühlenweg 1, 07743 Jena, Germany

**Keywords:** *Staphylococcus aureus*, pneumonia, aging, Terc, T cells, inflammation, macrophages

## Abstract

The telomerase RNA component (Terc) constitutes a non-coding RNA critical for telomerase function, commonly associated with aging and pivotal in immunomodulation during inflammation.

Our study unveils heightened susceptibility to pneumonia caused by *Staphylococcus aureus (S. aureus)* in *Terc* knockout (Terc^ko/ko^) mice compared to both young and old infected counterparts. The exacerbated infection in Terc^ko/ko^ mice correlates with heightened inflammation, manifested by elevated interleukin-1β (IL-1β) levels and activation of the NLR Family Pyrin Domain Containing 3 (NLRP3) inflammasome within the lung.

Employing mRNA sequencing methods alongside *in vitro* analysis of alveolar macrophages (AMs) and T cells, our study elucidates a compelling correlation between Terc^ko/ko^, inflammation, and impaired T cell functionality. *Terc* deletion results in compromised T cell function, characterized by dysregulation of the T cell receptor and absence of CD247, potentially compromising the host’s capacity to mount an effective immune response against *S. aureus*.

This investigation provides insights into the intricate mechanisms governing increased vulnerability to severe pneumonia in the context of Terc deficiency, which might also contribute to aging-related pathologies, while also revealing for the first time the influence of Terc on T cell function.

**Graphical abstract:** 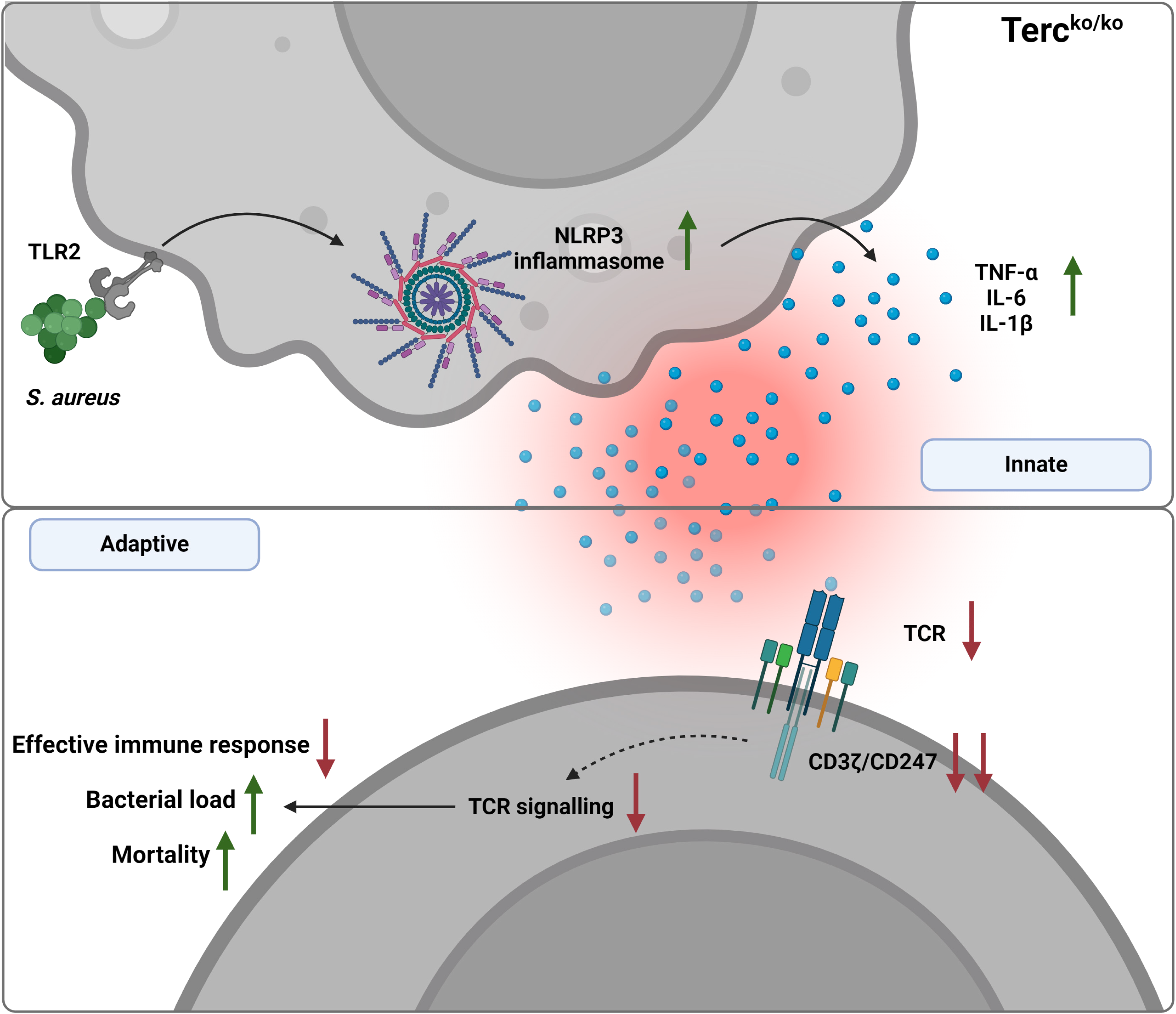

## Introduction

As the global population ages, the impact of infectious diseases on the aging population becomes increasingly significant. Among the most relevant risk factors leading to higher susceptibility as well as severity in both community-acquired (CA) and healthcare-associated (HA)-pneumonia is advanced age (≥ 60 years) (Torres et al., 2021). Particularly severe is pneumonia caused by *Staphylococcus aureus* (*S. aureus)* (Lee et al., 2022). *S. aureus* is a gram-positive bacterium that frequently colonizes humans but can manifest various pathologies, ranging from mild to more severe infections, including pneumonia (Howden et al., 2023; Jong, Kessel, & Strijp, 2019). It accounts for a notable proportion of CA and HA cases (Lee et al., 2022; Torres et al., 2021). The bacterium exhibits a broad spectrum of virulence factors, including toxins, adhesins, and immune evasion strategies, contributing to its ability to establish infections in the lower respiratory tract (Howden et al., 2023). Aging is a central factor predisposing individuals to a higher risk for *S. aureus* pneumonia (Torres et al., 2021), as aging is impacting several important functions of the body such as the immune response and thus the capability to mount an effective defensive response (Montecino-Rodriguez, Berent-Maoz, & Dorshkind, 2013). Age-related dysregulation of the immune response is also characterized by inflammaging, defined as the presence of elevated levels of pro-inflammatory cytokines in the absence of an obvious inflammatory trigger (Franceschi et al., 2000; Mogilenko, Shchukina, & Artyomov, 2022). Additionally, immune cells, such as macrophages, exhibit an activated state that alters their response to infection (Canan et al., 2014). In contrast, the immune response of macrophages to infectious challenges has been shown to be initially impaired in aged mice (Boe, Boule, & Kovacs, 2017). Thus aging is a relevant factor impacting the pulmonary immune response. This highlights the vital connection between aging and *S. aureus* pneumonia, as well as the need for further research investigating this connection.

The process of aging is a multifaceted phenomenon that involves various underlying mechanisms, known as hallmarks (López-Otín, Blasco, Partridge, Serrano, & Kroemer, 2023). Among these hallmarks, telomere shortening is a significant factor in cellular aging. Telomeres serve as protective caps at the end of chromosomes, which shorten with every cell division until they reach a critical length where replication becomes impossible, eventually resulting in either apoptosis or senescence (López-Otín et al., 2023). However, this process can be counteracted by the telomerase, which can add new repeats to the ends of chromosomes and thus prevent the shortening of telomeres (Greider & Blackburn, 1985). The telomerase mainly consists of two parts: the telomerase reverse transcriptase (Tert), which harbors the enzymatic activity, and Terc, which serves as a template for the addition of new repeats (Sahin & DePinho, 2010; Shay & Wright, 2019). Telomerase activity is restricted to specific tissues and cell types, largely dependent on the expression of *Tert*. While *Tert* is highly expressed in stem cells, progenitor cells, and germline cells, its expression is minimal in most differentiated cells (Chakravarti, LaBella, & DePinho, 2021). Consequently, the impact of telomerase dysfunction on tissues varies according to their self-renewal rate (Chakravarti et al., 2021).

One important aspect of telomere dysfunction is the impact of telomere shortening on the immune system as well as the hematopoietic system. Tissues or organ systems that are highly replicative, such as the skin or the hematopoietic system, are affected first by telomere shortening (Chakravarti et al., 2021). Due to the loss of the protective telomere caps, mutations can accumulate, and the cells ultimately become senescent (Chakravarti et al., 2021). Interestingly, while most differentiated cells do not exhibit telomerase activity, T cells display a high activity of this enzyme (Hodes, Hathcock, & Weng, 2002). Aging T cells become senescent, which is characterized by short telomeres and the absence of CD28 expression (Hohensinner, Goronzy, & Weyand, 2011). Furthermore, telomere length and T cell functionality are closely connected. T cells with shorter telomeres are implicated in diseases related to aging like arthritis (Hohensinner et al., 2011). Additionally, CD4^+^ T cells from Terc knockout (Terc^ko/ko^) mice displayed several characteristics of aged T cells such as a reduction in CD28 expression and changed secretion of cytokines. Furthermore the naïve T cell population was found to be reduced (Matthe, Thoma, Sperka, Neurath, & Waldner, 2022).This highlights the importance of intact telomeres and telomerase on immune cell function.

While most differentiated cells lack the expression of *Tert*, almost all human cells and several murine tissues express *Terc* (Zhang et al., 2018). However, the function of ubiquitous *Terc* expression remains elusive (Chakravarti et al., 2021; Shay & Wright, 2019). Interestingly, recent studies identified a variety of functions of Terc, which are independent of its function as part of the telomerase. Notably, a study could show that Terc triggers inflammation via the nuclear factor kappa-light-chain-enhancer of activated B cells (NF-κB) pathway (Liu, Yang, Ge, Liu, & Zhao, 2019). Furthermore, Terc was identified to be involved in cell proliferation by enhancing the expression of several genes belonging to the Phosphoinositide 3-kinase (Pi3K) – Protein Kinase B (AKT) pathway (Wu et al., 2022). The same study could show that this mechanism also contributed to CD4^+^ T cell expansion (Wu et al., 2022). Additionally, *Terc* deletion in mice induced NLR Family Pyrin Domain Containing 3 (NLRP3) inflammasome activation in macrophages upon pulmonary infection with *S. aureus*, thereby revealing another connection between immune modulation and the presence of Terc (Kang et al., 2018). Thus, it appears that Terc could serve as a relevant immunomodulatory factor, potentially implicated in *S. aureus-*induced pathologies. Interestingly, downregulation of *Terc* and *Tert* expression in tissues of aged mice and rats has been found (Tarry-Adkins, Aiken, Dearden, Fernandez-Twinn, & Ozanne, 2021; Zhang et al., 2018). Therefore, as a potential immunomodulatory factor reduced *Terc* expression could be connected to age-related pathologies.

Knockout of *Terc* in mice is a well-established model for premature aging, as telomeres of Terc^ko/ko^ mice shorten with every generation (Wong et al., 2008). Additionally, the telomeres shorten during each generation with increasing age of the mice (Wong et al., 2008). Furthermore, third-generation (G3) Terc^ko/ko^ mice with a C56Bl/6 background have a shorter lifespan, a smaller litter size and a generally reduced body size and weight (Wong et al., 2008). Although G3 Terc^ko/ko^ mice with shortened telomeres were used in this study, they were infected at a young age (8 weeks). This approach allowed for the investigation of *Terc* deletion effects rather than telomere dysfunction. As control cohort age-matched young WT mice were utilized. To investigate whether *Terc* deletion, beyond critical telomere shortening, impacts the pulmonary immune response, we used young Terc^ko/ko^ mice. Additionally, naturally aged mice (2 years old) were infected to explore the potential link to a fully developed aging phenotype.

In this study, we aimed to investigate the impact of *Terc* deletion, as an essential immunomodulatory factor, on the immune response to *S. aureus-*induced pneumonia in mice. For this purpose, G3 Terc^ko/ko^ mice were infected with *S. aureus*, and the progression of disease and immune response were evaluated. Strikingly, our data highlights that the deletion of *Terc* resulted in a more severe disease outcome and disrupted the innate and adaptive immune response. These findings suggest a possible connection between Terc and immune cell homeostasis.

## Materials and Methods

### Bacteria

All *in vivo* and *in vitro* experiments were carried out with the *S. aureus* strain USA300. *S. aureus* was cultivated in brain heart infusion (BHI) medium (Thermo Fisher Scientific, Waltham, MA, US) at 37°C while shaking at 150 rpm. The bacteria were harvested during the mid-logarithmic phase and adjusted to the appropriate optical density (OD) at 600nm for the respective experiments in phosphate-buffered saline (PBS). The bacteria were then either used freshly for *in vivo* infection of mice or stored at −80 °C until further usage in *in vitro* experiments.

For heat-killing, the adjusted bacteria stock was incubated at 95°C for 5 minutes while shaking at 500 rpm. A fraction of the heat-killed bacteria was plated on Müller-Hinton (MH) plates. Plates were incubated for 5 days at 37°C to confirm heat-killing of the bacteria.

### Determination of bacterial load

Organs were homogenized in the appropriate amount of PBS (adjusted to their weight in mg) using a SpeedMill Plus (Analytik Jena, Jena, Germany). Homogenized organs and bronchoalveolar lavage (BAL) fluid were diluted in a decadic dilution series and plated onto MH plates. Plates were incubated overnight at 37°C. Colonies were counted, and colony-forming units (CFU) per ml was calculated.

### Mouse models and infection

Mouse *in vivo* experiments were approved by the Office for Consumer Protection of Thuringia (TV-Number: UKJ-19-028 and UKJ-22-023).

G3 female Terc^ko/ko^ mice with a C57Bl/6 background were bred from homozygous parents in the animal facility of the Jena University Hospital. To confirm knockout of *Terc*, DNA was extracted from the tail of newborn mice using the DNeasy Blood & Tissue Kit (Qiagen, Venlo, Netherlands) according to the manufacturer’s protocol. The DNA was then used for genotyping via PCR using the following primer sequences: mTR-R: 5’-TTC TGA CCA CCA CCA ACT TCA AT-3’; mTR-WT-F: 5’-CTA AGC CGG CAC TCC TTA CAA G-3’; 5PPgK(-F): 5’-GGG GCT GCT AAA GCG CAT-3’. The results were analyzed with an Agilent 2200 TapeStation system using a Agilent D1000 ScreenTape (both Agilent Technologies, Santa Clara, CA, USA). The WT *Terc* product had a size of 200 bp while the knockout product had a size of 180 bp. Only mice that showed a single band at 180 bp were used for the further breeding process.

Terc^ko/ko^ mice aged 8 weeks, were used for infection studies (n = 8; non-infected = 3; infected = 5). Female young WT (age 8 weeks) and old WT (age 24 months) C57Bl/6 mice (both n = 10; non-infected = 5; infected = 5) were purchased from Janvier Labs (Le Genest-Saint-Isle, France). All infected mouse cohorts were compared to their respective non-infected controls, as well as to the infected groups from other cohorts. Additionally, comparisons were made between the non-infected cohorts across all groups.

All mice were anesthetized with 2% isoflurane before intranasal infection with *S. aureus* USA300 (1×10^8^ CFU/20µl) per mouse. After 24 hours, the mice were weighed and scored as previously described (Hornung et al., 2023). Infected Terc^ko/ko^ mice were grouped into different degrees of severity based on their clinical score, fatal outcome of the disease (fatal) and the presence of bacteria in organs other than the lung (systemic infection) for the indicated analysis. Mice with fatal infections were excluded from subsequent analyses, with only their final scores being reported. The mice were sacrificed via injection of an overdose of xylazine/ketamine and bleeding of axillary artery after 24 hpi. BAL was collected by instillation and subsequent retrieval of PBS into the lungs. Serum and organs were collected. Bacterial load in the BAL, kidney and liver was determined by plating of serially diluted sample as described above. For this organs were previously homogenized in the appropriate volume of PBS. Gene expression was analyzed in the right superior lung lobe. Lobes were therefore homogenized in the appropriate amount of TriZol LS reagent (Thermo Fisher Scientific, Waltham, MA, US) prior to RNA extraction. The left lung lobe was embedded into Tissue Tek O.C.T. (science services, Munich, Germany) and stored at −80°C until further processing for histological analysis. Cytokine measurements were performed using the right inferior lung lobe. Lobes were previously homogenized in the appropriate volume of PBS. Remaining organs were stored at −80°C until further usage.

Mouse studies were conducted without the use of randomization or blinding.

### Telomere length measurement

The length of telomeres in the lung of young WT, old WT and Terc^ko/ko^ mice was measured using the Absolute Mouse Telomere Length Quantification qPCR Assay Kit (ScienCell research Laboratories, Carlsbad, CA, USA) according to the manufacturer’s protocol. Genomic DNA from the lungs was extracted by DNeasy Blood & Tissue Kit (Qiagen, Venlo, Netherlands) according to the manufacturer’s protocol. For the PCR 2ng of each sample was used. Average telomere length per chromosome end was calculated as described by the manufacturer.

### Histology and Immunofluorescence

Preparation and staining of tissue cryosections was performed as described previously (Hornung et al., 2023).

For staining of *S. aureus,* rabbit anti-*S. aureus* polyclonal antibody (Cat. # PA1-7246, Thermo Fisher Scientific, Waltham, MA, US) was diluted 1:400 in antibody diluent with background reducing components (Agilent, Santa Clara, CA, US). Alexa Fluor® 488 AffiniPure Donkey Anti-Rabbit IgG (Cat. #711-545-152, Jackson Immuno Research, West Grove, PA, US) and BODIPY 558/568 phalloidin, which stains actin fibers, (Cat. # B3475, Thermo Fisher Scientific, Waltham, MA, US) were diluted in antibody diluent 1:500 and 1:400 respectively and applied to the slides as secondary antibodies.

### Immunocytochemistry

Murine primary macrophages were fixed with 4% paraformaldehyde (PFA) for 15 minutes at 37°C, subsequently permeabilized with 0.1% Triton-X, and blocked with 3% bovine serum albumin (BSA) in PBS for each 30 minutes at room temperature (RT). Cells were incubated overnight with either rabbit anti-*S. aureus* polyclonal antibody or rabbit anti-CD68 polyclonal antibody (Cat. # BOSSBS-0649R, VWR, Radnor, PA, US) diluted 1:400 or 1:200 in 3% BSA in PBS respectively. The following day, the slides were washed three times with PBS. Cy3-conjugated AffiniPure Donkey Anti-Rabbit IgG (Cat. #711-165-152, Jackson Immuno Research, West Grove, PA, US) and Alexa Fluor™Plus 647 Phalloidin (Cat. # A30107, Thermo Fisher Scientific, Waltham, MA, US) were applied to the cells diluted 1:500 and 1:400 in 3% BSA in PBS respectively and incubated for 1 hour at RT. The cells were then mounted with DAPI Fluoromount-G.

The immunofluorescence pictures were taken with an AxioObserver Z.1 microscope (Carl Zeiss AG, Oberkochen, Germany) and analyzed with the Zen software (Zen Pro v3.3).

### RNA extraction and mRNA sequencing

RNA was extracted from murine lung tissue using the Trizol/Chloroform method. First, about 15 mg of lung tissue was homogenized in TriZol LS reagent (Thermo Fisher Scientific, Waltham, MA, US) using the SpeedMill Plus. The tissue-free supernatant of the homogenate was transferred into a new tube after centrifugation. 50 µl of chloroform was added to 250 µl TriZol Reagent, incubated for 3 minutes at RT, and centrifuged at 12000xg for 15 minutes at 4°C. The aqueous phase was mixed with isopropanol in a 1:2 ratio and incubated at RT for 10 minutes before centrifugation at 12000xg for 10 minutes at 4°C. Afterward, the RNA pellet was washed twice with 75% ethanol. Finally, ethanol was removed, and the pellet was air-dried for 5-10 minutes. The dry pellet was then dissolved in sterile distilled water, and RNA concentration was determined by an ND-1000 NanoDrop spectrophotometer (PEQLAB Biotechnologie GmbH, Erlangen, Germany). Before sequencing, RNA integrity was measured using a 5400 Fragment Analyzer (Agilent Technologies, Santa Clara, CA, US). RNA concentration and RNA integrity of the samples can be found in Supplemental Table 1.

Library construction and mRNA sequencing were performed by Novogene Co., LTD. (Beijing, China) using the Illumina platform Novaseq 6000 S4 flowcell V1.0, based on the sequencing by synthesis (SBS) mechanism and PE150 strategy. For bioinformatics analysis, raw reads in FASTQ format were first processed using fastp. Reads containing adapters and poly N sequences were removed as well as low quality reads to generate clean data. Clean reads were then mapped to the reference genome (Mus Musculus (GRCm39/mm39)) with the HISAT2 software. Gene expression levels were quantified using the FPKM method. DESeq2 software as well as negative binomial distribution was used for analysis of differential gene expression. For FDR (False discovery rate) calculation the Benjamini-Hochberg procedure was used. A summary of the quality of the sequencing data of each sample can be found in Supplemental Table 2.

Heatmaps of the differentially regulated genes (DEG) were constructed using R (v4.2.2; R Foundation for Statistical Computing, https://www.r-project.org/). Only significant DEGs (p-value < 0.05) were displayed in the heatmaps. Non-significant genes were set to zero and are shown in white. All DEGs used for construction of the respective heatmaps are summarized in Supplemental Table 3a-i.

### Immunophenotyping of the spleen

Mice were sacrificed as described above, and spleens were freshly stored in cold 2% FCS in PBS. For isolation of immune cells, spleens were mashed through a 70 µm cell strainer (Miltenyi Biotec, Bergisch Gladbach, Germany), and red blood cell lysis was performed by adding 5 ml of Ammonium–chloride–potassium (ACK) lysis buffer (Thermo Fisher Scientific, Waltham, MA, US) for 5 minutes at 4°C to the pelleted cells. Cells were then counted, adjusted to 2×10^6^ per ml PBS, and stained with BD Fixable Viability Stain 780 (BD Biosciences, Heidelberg, Germany) diluted 1:1000 in PBS for 15 minutes at RT. Non-specific binding of antibodies was blocked by incubating the samples with 3% rat serum in BD staining buffer (BD Biosciences, Heidelberg, Germany) for 5 minutes at RT. Anti-CD11c (Cat. #557401), anti-CD19 (Cat. #566412), anti-CD335 (Cat. #564069), anti-CD34 (Cat. #742971), anti-CD3ε (Cat. #553066), anti-Ly6G (Cat. #560601), and anti-F4/80 (Cat. #565411) were added to the samples in a dilution of 1:100 and incubated for 20 minutes at 4°C. All antibodies were purchased from BD Biosciences. Stained samples were measured with a BD Symphony A1 (BD Biosciences, Heidelberg, Germany), and analysis was performed with FlowJo v10.8.1. Detailed information on the gating strategy is presented in Supplemental Figure 3A.

### Isolation and infection of murine T cells and alveolar macrophages

Terc^ko/ko^ and young WT mice were sacrificed. Killing for scientific purposes was conducted in accordance with the German animal welfare regulations. Alveolar macrophages (AMs) were isolated as published elsewhere (Rios, Touyz, & Montezano, 2017). In short, the murine lungs were washed out several times with PBS containing 0.5 mM Ethylenediaminetetraacetic acid (EDTA). The lavage was centrifuged at 350xg for 10 minutes at 4°C. The cell pellet was dissolved in Roswell Park Memorial Institute (RPMI) medium containing 20 mM HEPES and 5% fetal calf serum (FCS) and counted. Subsequently, cells were seeded and incubated at 37°C for 2 hours.

Lymphocytes were isolated from the spleen as described above. To subsequently isolate CD4^+^ T cells, cells were separated via magnetic cell separation. These cells were mixed with CD4^+^ beads and separated via MACS LS Columns (both Miltenyi Biotec, Bergisch Gladbach, Germany) according to the manufacturer’s protocol. The purity of the resulting CD4^+^ T cell population was measured via flow cytometry using a BD Symphony A1 (BD Biosciences, Heidelberg, Germany). T cells were added to AMs for co-culture in a 1:1 ratio or cultured alone.

For infection with heat-killed (HK) *S. aureus,* a multiplicity of infection (MOI) of 5 was calculated, and the appropriate bacteria volume was added to each well. Cells were then incubated for 24 hours and harvested for downstream analysis.

### Cytokine measurement

For cytokine measurements, the right inferior lung lobe was homogenized using a SpeedMill Plus in the appropriate volume of PBS to adjust the organs to their weight in mg. Then, the cytokines were measured with the LEGENDPlex^TM^ mouse inflammation panel (BioLegend, San Diego, CA, US) according to the manufacturer’s instructions. Samples were run in duplicates on the same plate. For the young WT and Terc^ko/ko^ cohort three biological replicates and for old WT cohorts four biological replicates were measured. Supernatants of co-cultured and single-cultured T cells were measured with the LEGENDPlex^TM^ mouse T helper cytokine panel (BioLegend, San Diego, CA, US). For each condition three biological replicates were measured. All samples were run on the same plate. Flow cytometry analysis was performed with a BD Symphony A1, and data analysis was carried out with the Qognit software v2023-02-15 (BD Biosciences, Heidelberg, Germany).

### Flow cytometry

Co-cultured and single-cultured T cells were transferred to FACS tubes and centrifuged at 500xg for 5 minutes at 4°C. The supernatant was frozen for cytokine measurements, and the cell pellet was stained with PBS containing 1:1000 diluted BD Fixable Viability Stain 780 (Cat. # 565388, BD Biosciences, Heidelberg, Germany) for 15 minutes at RT. Staining of surface markers was performed as described above. Samples were stained with anti-CD3ε (1:100, Cat. #553066), anti-CD25 (1:800, Cat. #553866), anti-CD69 (1:400, Cat. # 551113) and anti-CD44 (1:100, Cat. # 560569) for 30 minutes at 4°C. All antibodies were purchased from BD Pharmingen. Subsequently, samples were permeabilized and fixed. For this, the samples were centrifuged for 5 minutes at 500xg and 4°C, resuspended in BD Cytofix/Cytoperm buffer (BD Biosciences, Heidelberg, Germany), and incubated for 20 minutes at 4°C. Samples were washed once with BD Perm/Wash Buffer (BD Biosciences, Heidelberg, Germany). For intracellular staining, samples were stained with anti-CD247 (1:100, Cat. # ab91493, Abcam, Cambridge, UK) antibody in BD Perm/Wash Buffer for 30 minutes at 4°C. Lastly, the samples were washed and resuspended in BD Perm/Wash Buffer. Stained samples were measured with a BD Symphony A1 and analyzed with FlowJo v10.8.1. Detailed information on the gating strategy is presented in Supplemental Figure 4B.

### Scanning electron microscopy (SEM)

T cells and AMs were isolated as described above. Cells were grown on poly-L-lysine (Merck-Millipore, Burlington, MA, US) coated glass coverslips in a 1:1 ratio. T cells were allowed to attach to the coverslips for 30 minutes before adding *S. aureus*. After adhesion of T cells, *S. aureus* USA300 was added at a MOI1 and incubated at 37°C. After 3.5 hours, the samples were fixed with freshly prepared modified Karnovsky fixative (4% w/v paraformaldehyde, 2.5 % v/v glutaraldehyde in 0.1 M sodium cacodylate buffer, pH 7.4) for 1 hour at RT. After washing each 3 times for 15 minutes with 0.1 M sodium cacodylate buffer (pH 7.4), the cells were post-fixed with 2% (w/v) osmium tetroxide for 1 hour at RT. Subsequently, the samples were dehydrated in ascending ethanol concentrations (30, 50, 70, 90, and 100%) for 15 minutes each. Next, the samples were critical-point dried using liquid CO_2_ and sputter coated with gold (thickness approx. 2 nm) using a CCU-010 sputter coater (safematic GmbH, Zizers, Switzerland). Finally, the specimens were investigated with a field emission SEM LEO-1530 Gemini (Carl Zeiss NTS GmbH, Oberkochen, Germany).

### Statistical analysis and scheme design

Data are presented as mean ± SD, or as median with interquartile range for violin and box plots, with up to four levels of statistical significance indicated. P-values were calculated using Kruskal-Wallis test. Individual replicates are represented as single data points. Statistical analysis was carried out using GraphPad Prism v 9.0. Illustrations were created with BioRender.com.

## Results

### *Terc* knockout is associated with severe pneumonia in mice

To investigate the impact of Terc on the course of *S. aureus* pneumonia, we compared young Terc^ko/ko^ mice to young and old wild type (WT) mice. For this purpose, we performed an intranasal infection of Terc^ko/ko^ mice at the age of 8 weeks and naturally aged mice at the age of 2 years (old WT) with the *S. aureus* strain USA300. WT mice at the age of 8 weeks (young WT) were additionally infected and utilized as the control group (Figure 1A).

**Figure 1:**
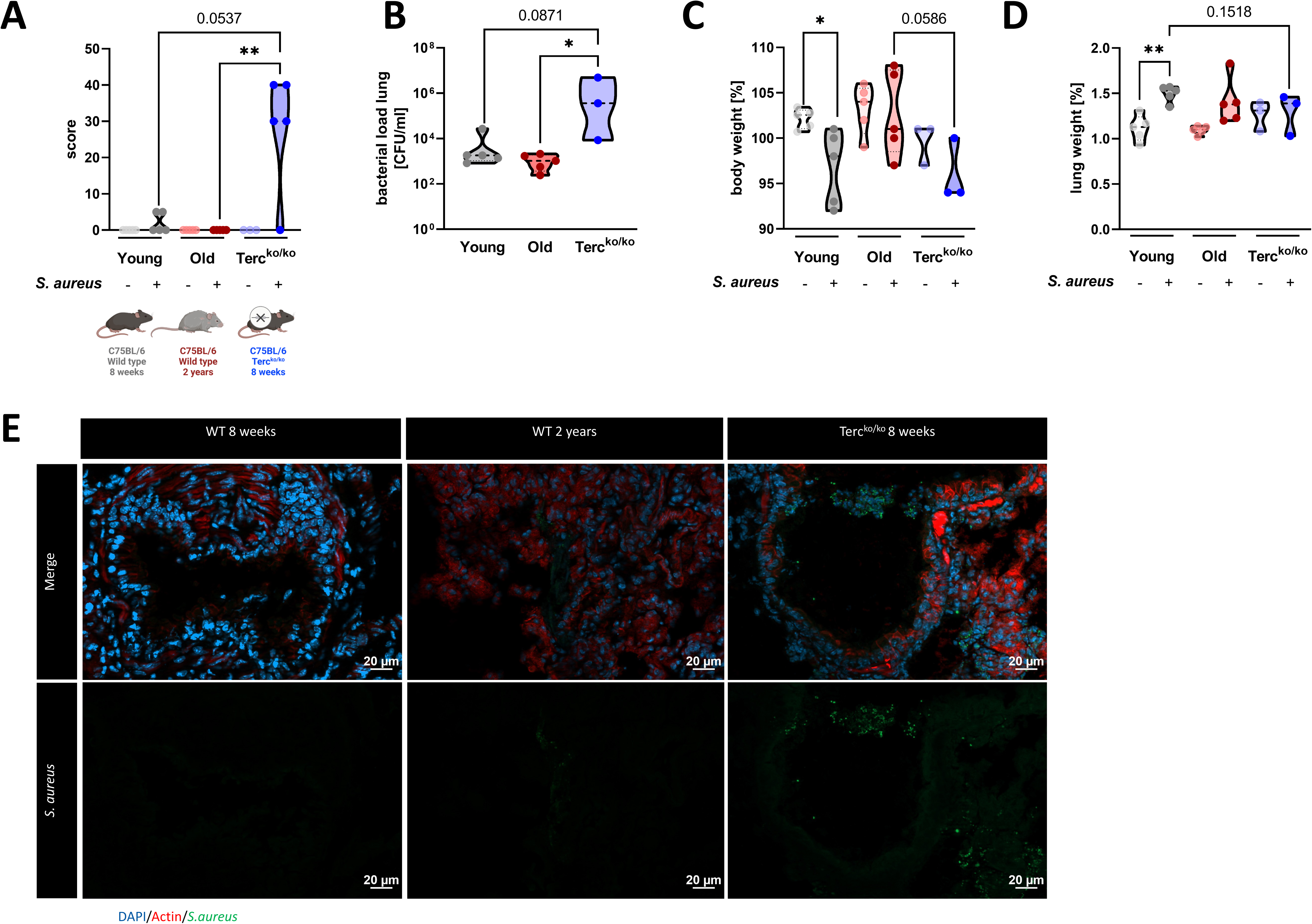
Terc^ko/ko^ mice exhibit more severe pneumonia with increased mortality and a higher bacterial load. A) Clinical score of non-infected and infected young WT (n =10, 5 non-infected and 5 infected), old WT (n =10, 5 non-infected and 5 infected), and Terc^ko/ko^ (n =8, 3 non-infected and 5 infected) mice at 24 hpi. The age and genetic background of the different groups is depicted below. B) Bacterial load of the lungs of infected mice at 24 hpi. Data is displayed as logarithmic. C) Relative body weight of young WT, old WT and Terc^ko/ko^ mice. Relative body weight displays body weight at 24 hpi as a percentage of the body weight at the time of infection. D) Relative lung weight of young WT, old WT, and Terc^ko/ko^ mice. Relative lung weight displays lung weight at 24 hpi as a percentage of the current body weight. E) Immunofluorescence staining of lung tissue of infected young (WT 8 weeks), old (WT 2 years), and Terc^ko/ko^ mice. Lungs were stained for *S. aureus* (green), actin (red), and DAPI (blue). Representative pictures are shown for each group. *p < 0.05, **p < 0.01, ***p<0.001. P calculated by Kruskal-Wallis-test (A-D). Data is displayed as violin plot showing the median as well as lower and upper percentile of each dataset. Each replicate is displayed as a data point.

The clinical score of infected Terc^ko/ko^ mice was increased compared to the other two infected mice cohorts at 24 hours post infection (hpi) (Figure 1A). The bacterial load was significantly higher in the BAL of Terc^ko/ko^ mice compared to old WT mice (Figure 1B). Interestingly, three distinct degrees of severity were observed in the infected Terc^ko/ko^ mice (n = 5): mild infection, systemic infections, or fatal infections (Supplemental Figure 1A). Mice exhibiting systemic infection were characterized by the presence of bacteria in extra pulmonary organs, specifically observed in the liver and kidney tissues of some Terc^ko/ko^ mice (Supplemental Figure 1B).

Infection of young WT mice resulted in a significant decrease in bodyweight at 24 hpi. The body weight of infected Terc^ko/ko^ mice was reduced compared to the infected old WT mice and the non-infected Terc^ko/ko^ mice (Figure 1C). In contrast, at 24 hpi the relative weight of the lung was increased in infected compared to the non-infected mice for all cohorts, indicating inflammation of the lung (Figure 1D). However, this effect was the most pronounced in young WT mice.

Based on our observation of the different degrees of severity displayed by the infected Terc^ko/ko^ mice, we categorized them according to their clinical score and the presence of bacteria in extra pulmonary organs, distinguishing between mice with and without systemic infection. This grouping resulted in more pronounced differences between the mice cohorts regarding bacterial load and reduction of relative body weight (Supplemental Figure 1C-D). The relative lung weight increased for all groups compared to the respective non-infected mice, except for the Terc^ko/ko^ mouse without systemic infection (Supplemental Figure 1E). Additionally, average telomere length of the lungs of young and old WT as well as Terc^ko/ko^ mice was measured. As shortened telomeres are a hallmark of aging, impacting other cellular mechanisms such as cellular senescence, they could have an influence on the disease progression (López-Otín et al., 2023). Notably, the telomeres of Terc^ko/ko^ mice were the shortest among the three different mouse models, even shorter than those of naturally aged mice (Supplemental Figure 1F).

In order to investigate gene expression patterns, mRNA sequencing of the lungs was performed. This data supported the grouping of infected Terc^ko/ko^ mice into different groups, as the principal component analysis (PCA) plot showed a separate cluster containing Terc^ko/ko^ mice with systemic infection (Supplemental Figure 1G).

Immunofluorescence (IF) staining of *S. aureus* within the lung tissue of all three infected mice cohorts was carried out. The increased signal for *S. aureus* in sections of Terc^ko/ko^ mice compared to sections of young WT and old WT mice closely reflects the quantified CFU detected in the BAL (Figure 1E).

Terc^ko/ko^ mice were characterized by an increased bacterial load in the lungs, a higher clinical score as well as mortality (Figure 1A-B; Supplemental Figure 1A). Additionally, we could observe different degrees of severity in the infected Terc^ko/ko^ cohort and could identify a subgroup of mice with systemic infection (Supplemental Figure 1A-B and G).

### Terc^ko/ko^ mice showed excessive inflammation with inflammasome activation

In order to investigate the underlying pathomechanisms leading to a more severe course of disease in the Terc^ko/ko^ mice, inflammation parameters in the lungs were measured. In the course of this analysis, we applied the previously outlined groups of mice, differentiating between those with and without systemic infection in the Terc^ko/ko^ cohort. Several pro-inflammatory cytokines were significantly upregulated in Terc^ko/ko^ mice with systemic infection, such as Interleukin-6 (IL-6), Granulocyte-macrophage colony-stimulating factor (GM-CSF), and interleukin-1β (IL-1β). Other important pro-inflammatory factors like Tumor necrosis factor α (TNFα) and Interleukin-1α (IL-1α) were also elevated (Figure 2A). A significant upregulation of pro-inflammatory mediators due to infection was, however; solely detected in infected Terc^ko/ko^ mice compared to the corresponding non-infected control cohort (Figure 2A).

**Figure 2:**
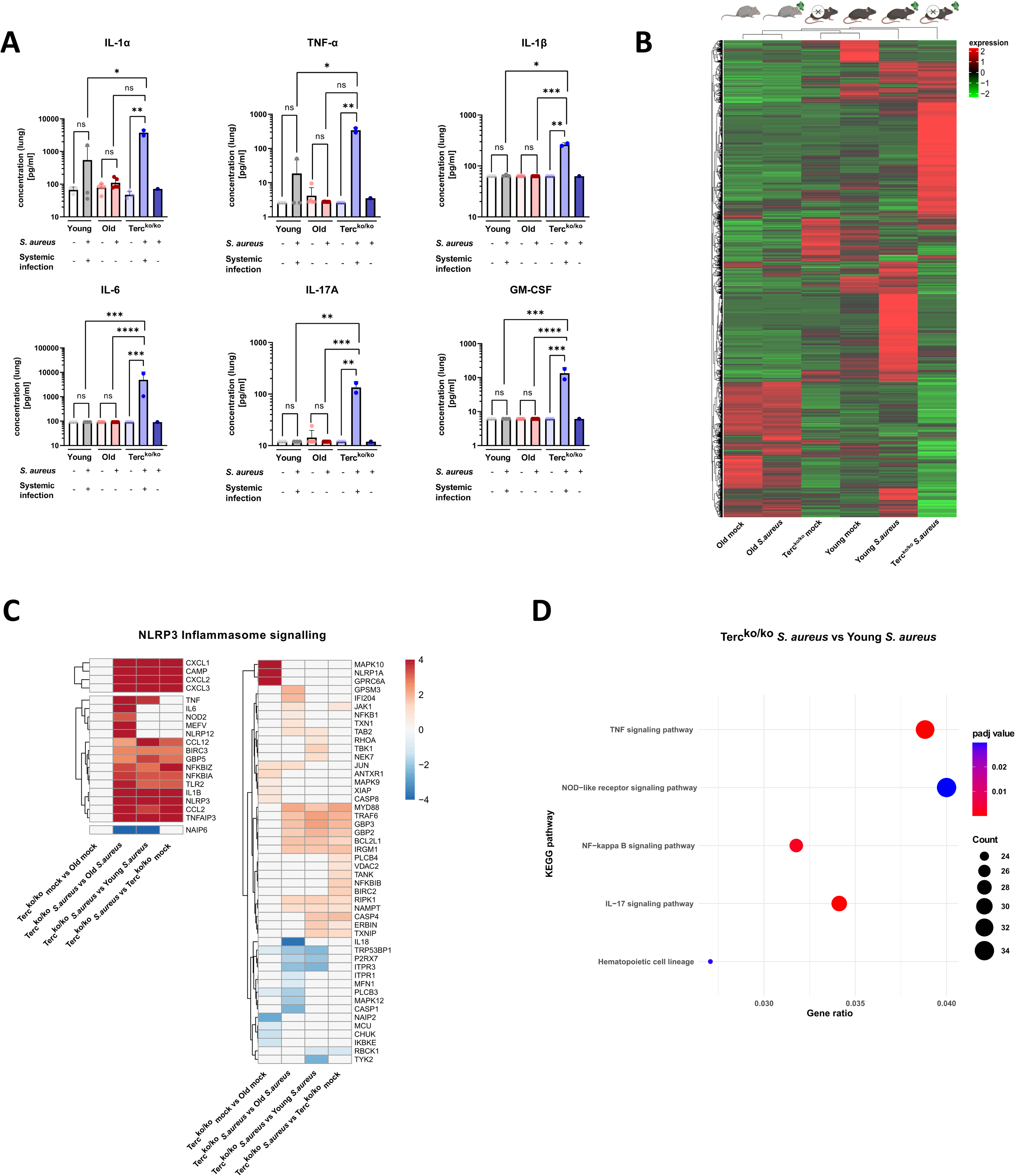
Lungs of Terc^ko/ko^ mice display excessive inflammation and NLRP3 inflammasome activation. A) Pro-inflammatory cytokines measured in lung homogenates of non-infected and infected young WT (each n=3), old WT (each n=4) and Terc^ko/ko^ mice (non-infected: n=3, with systemic infection: n=2 and without systemic infection: n=1). Infected Terc^ko/ko^ mice were grouped into mice with systemic and without systemic infection. Cytokines were measured using a flow cytometry based LEGENDPlex^TM^ mouse inflammation panel. Data is displayed on a logarithmic scale. B) Heatmap of overall gene expression in the lungs of the different mice cohorts at 24 hpi. Gene expression is displayed as the log2 of the fragments per kilobase of transcript per million fragments mapped (fpkm) of each individual gene. Gene expression is normalized to sequencing depth and transcript length. Red indicates upregulation, and green indicates downregulation of the gene expression. The respective mouse groups are visualized above the heatmap. For each mouse cohort three biological replicates were sequenced. C) Heatmap of differentially expressed genes (DEG) belonging to the NLRP3 inflammasome pathway. Red indicates upregulation, and blue indicates downregulation of the gene. Differential gene expression analysis was conducted using the DESeq2 software as well as negative binomial distribution. For FDR calculation the Benjamini-Hochberg procedure was used. Heatmaps were constructed using R Studio. Only significant DEGs (p-value < 0.05) were displayed in the heatmap. Non-significant genes were set to zero and are shown as white. DEGs are displayed as log2fold-change. For each group three biological replicates were sequenced and used for analysis. For clarity, the heatmap has been split in half, and no column clustering has been performed. The respective groups used for the comparison are indicated below each column. D) KEGG pathway enrichment analysis comparing infected young and Terc^ko/ko^ mice at 24 hpi. The gene ratio describes the ratio of differentially expressed genes to all genes of the respective pathway. The dot plot was constructed using R Studio. All sequencing data displays mRNA sequencing data of non-infected and infected lungs of the three different mice cohorts at 24 hpi. ns ≥ 0.05, *p < 0.05, **p < 0.01, ***p<0.001, ****p < 0.0001. P calculated by Kruskal-Wallis-test (A). Data is presented as mean ±SD. Each replicate is displayed as a data point.

We performed mRNA sequencing of the murine lung tissue of infected and non-infected mice at 24 hpi to elucidate potential differentially expressed genes that contribute to the more severe illness of Terc^ko/ko^ mice. Comparison of overall gene expression pattern of the different groups revealed that non-infected young WT and Terc^ko/ko^ mice cluster together and show a similar expression pattern. In contrast, the infected mice of those cohorts display different expression patterns (Figure 2B). This highlights that most differences between the knockout and WT mice are only present after infection with *S. aureus*. Furthermore, both non-infected and infected old WT mice show a completely distinct expression profile compared to young WT as well as Terc^ko/ko^ mice (Figure 2B).

Intriguingly, several genes related to the NLRP3 inflammasome pathway, such as *NLRP3* or Thioredoxin Interacting Protein *(TXNIP)*, were upregulated in infected Terc^ko/ko^, indicating activation of the inflammasome (Figure 2C; Supplemental Figure 2A). Notably, expression of genes encoding for TNFα, IL-1β, and IL-6 were heightened. Thus, the increased inflammation in the lungs of Terc^ko/ko^ mice is likely induced by activation of the NLRP3 inflammasome (Figure 2C; Supplemental Figure 2A). Supporting this, the Nucleotide oligomerization domain (NOD)-like receptor signaling pathway was among the most significantly enriched pathways when comparing infected Terc^ko/ko^ and young WT mice (Figure 2D).

Macrophages are part of the first line of defense against bacterial infections as they can phagocytose bacteria and support pathogen clearance. AMs are present in the lung and can directly react to the pathogen at the site of infection via the production of inflammatory mediators or attraction of other immune cells (Malainou, Abdin, Lachmann, Matt, & Herold, 2023). To specifically investigate the role of AMs during *S. aureus* infection, we isolated them from lungs of young WT and Terc^ko/ko^ mice. By using scanning electron microscopy (SEM) we could strikingly visualize the ability of AMs to phagocytose *S. aureus* (Figure 3A). Additionally, we identified the AMs via IF staining of the macrophage marker CD68 (Supplemental Figure 2B).

**Figure 3:**
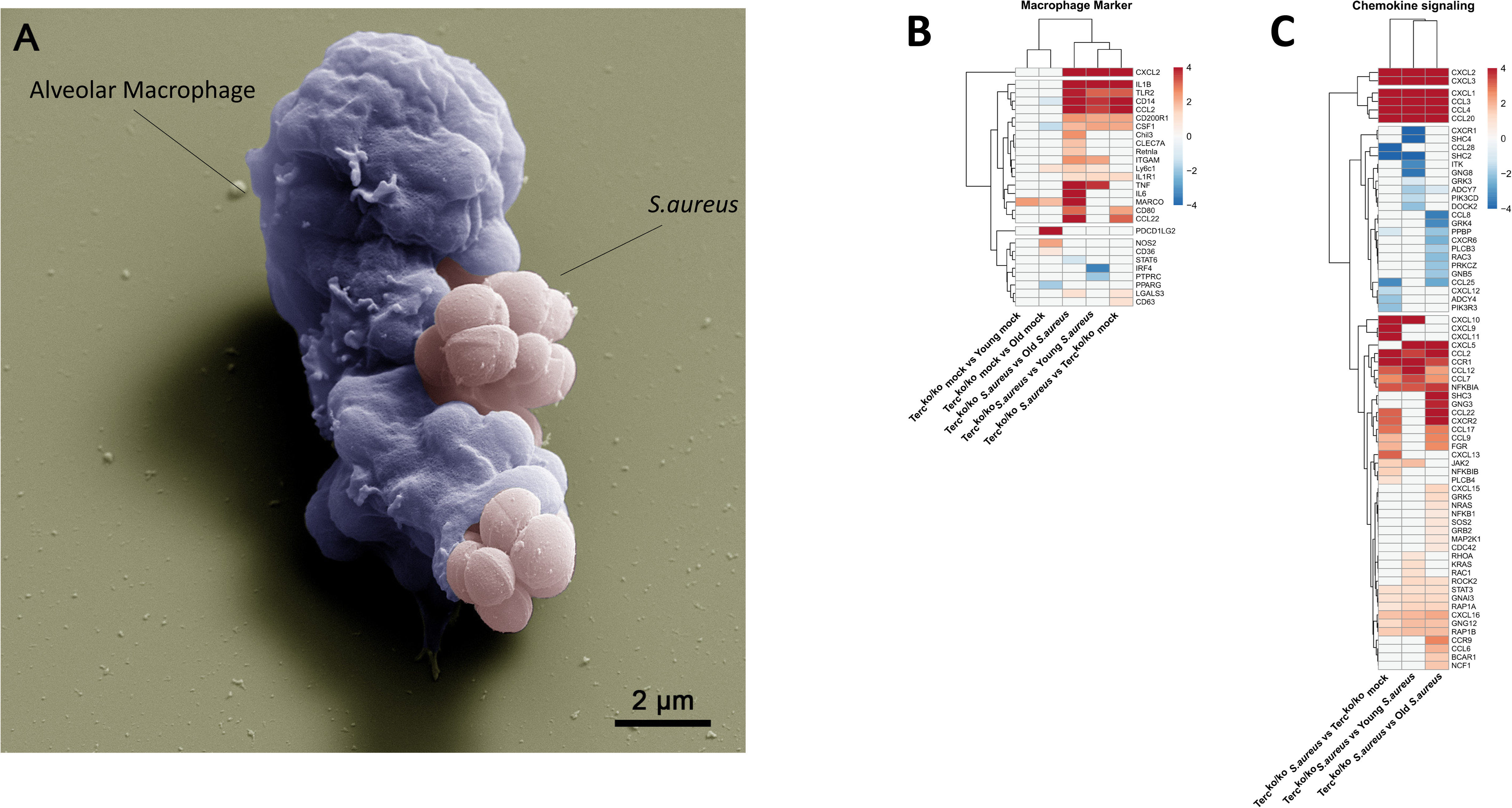
Elevated expression of pro-inflammatory M1 macrophages markers and chemokines are present in the lungs of infected Terc^ko/ko^ mice. A) Colorized SEM picture of an AM isolated from young WT mice phagocytosing *S. aureus*. Heatmap of differentially expressed macrophage markers (B) and DEGs belonging to the chemokine signaling pathway (C). Red indicates upregulation, and blue indicates downregulation of the gene. Differential gene expression analysis was conducted using the DESeq2 software as well as negative binomial distribution. For FDR calculation the Benjamini-Hochberg procedure was used. Heatmaps were constructed using R Studio. Only significant DEGs (p-value < 0.05) were displayed in the heatmap. Non-significant genes were set to zero and are shown as white. DEGs are displayed as log2fold-change. For each group three biological replicates were sequenced and used for analysis. The respective groups used for the comparison are indicated below each column. All heatmaps display mRNA sequencing data of non-infected and infected lungs of the three different mice cohorts at 24 hpi.

Notably, our sequencing data revealed an upregulation of pro-inflammatory M1 macrophage markers, such as C-X-C Motif Chemokine Ligand 2*(CXCL2)*, in Terc^ko/ko^ mice following infection (Figure 3B; Supplemental Figure 2C). Anti-inflammatory M2 markers, e.g., Interferon Regulatory Factor 4 *(IRF4)*, were mainly downregulated. Additionally, cluster of differentiation 14 *(CD14)*, a general macrophage marker, was increased in the infected Terc^ko/ko^ mice (Figure 3B; Supplemental Figure 2C). Furthermore, the expression of multiple chemokines such as C-X-C Motif Ligand 1-3 *(CXCL1-3)* and C-C Motif Chemokine Ligand 2 *(CCL2)* was elevated, suggesting increased migration of leukocytes such as macrophages and neutrophils to the lung (Figure 3C; Supplemental Figure 2D). Additionally, the TNF, NF-κB and IL-17 signaling pathways were among the most significantly enriched pathways in infected Terc^ko/ko^ mice compared to young WT mice (Figure 2D).

Combining the findings above, *S. aureus* infection in Terc^ko/ko^ mice leads to excessive inflammation induced by NLRP3 inflammasome activation and infiltration of pro-inflammatory immune cells, such as M1 macrophages.

### The T cell receptor is partially downregulated in infected Terc^ko/ko^ mice

In addition to macrophages, T cells and their activation play a crucial role in the immune response and are vital for resolving infections. Surprisingly, multiple general T cell markers such as *CD2*, *CD5*, and *CD4* were downregulated in the infected Terc^ko/ko^ mice compared to young WT mice (Figure 4A; Supplemental Figure 2E). However, these distinctions are not evident in comparisons between non-infected Terc^ko/ko^ and young WT mice. This further emphasizes that the observed variations in immune response activation arise specifically in response to infection (Figure 4A). Some T cell markers, such as *CD5*, that were downregulated in infected Terc^ko/ko^ mice were also downregulated in infected old WT mice compared to young WT mice (Supplemental Figure 2E).

**Figure 4:**
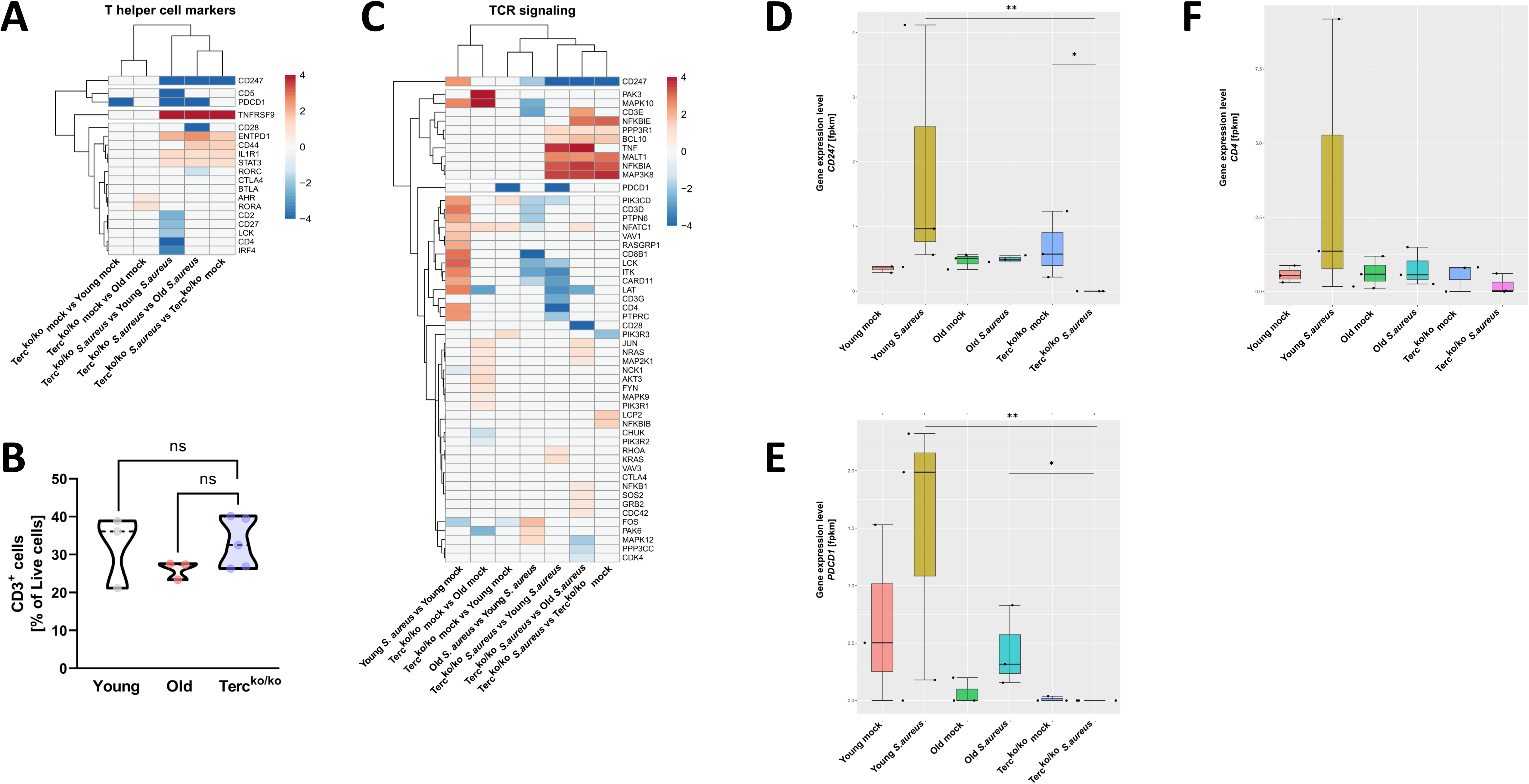
TCR in infected Terc^ko/ko^ mice is partially downregulated. Heatmap of differentially expressed T helper cell markers (A) and DEGs belonging to the TCR signaling pathway (C). Red indicates upregulation, and blue indicates downregulation of the gene. Differential gene expression analysis was conducted using the DESeq2 software as well as negative binomial distribution. For FDR calculation the Benjamini-Hochberg procedure was used. Heatmaps were constructed using R Studio. Only significant DEGs (p-value < 0.05) were displayed in the heatmap. Non-significant genes were set to zero and are shown as white. DEGs are displayed as log2fold-change. For each mouse cohort three biological replicates were sequenced and used for analysis. The respective groups used for the comparison are indicated below each column. All heatmaps display mRNA sequencing data of non-infected and infected lungs of the three different mice cohorts at 24 hpi. B) Percentage of CD3^+^ -living cells in non-infected young WT (n=3), old WT (n=3), and Terc^ko/ko^ mice (n=5) spleen. Cells were analyzed with flow cytometry. Gene expression levels in fpkm of *CD247* (D), *PDCD1* (E), and *CD4* (F) in the lungs of non-infected and infected young WT (n=3), old WT (n=3), and Terc^ko/ko^ mice (n=3) at 24 hpi. ns ≥ 0.05, *p < 0.05, **p < 0.01. P calculated by Kruskal-Wallis-test (B; D-F). Data is displayed as violin or boxplot showing the median as well as lower and upper percentile of each dataset. Each replicate is displayed as a data point.

To rule out that this effect is caused by a reduction of T cells in Terc^ko/ko^ mice, the immune cells of non-infected Terc^ko/ko^, young WT and old WT mice were analyzed via flow cytometry. However, no significant difference was observed between the groups (Figure 4B).

Additionally, several genes belonging to the T cell receptor (TCR) signaling pathway were downregulated in infected Terc^ko/ko^ mice, such as *CD3G* (encoding CD3γ) and *CD247* encoding CD3ζ (Figure 4C, Supplemental Figure 2F). Expression of downstream factors of the TCR like Lymphocyte Cell-Specific Protein-Tyrosine Kinase (*LCK*), linker of activation (*LAT*) and IL-2 inducible T cell kinase (*ITK*) was also reduced (Figure 4C, Supplemental Figure 2F). We also noted several of these distinctions, including the downregulation of *LCK* and *ITK* when comparing infected old WT and young WT mice (Figure 4C). Since there was no reduction of T cells in Terc^ko/ko^ mice, the change in TCR signaling and marker expression is likely induced by the infection.

Specifically, *CD247* was the only gene downregulated compared to infected young WT, old WT, and non-infected Terc^ko/ko^ mice (Figure 4C). Additionally, *CD247* was downregulated in infected old WT mice compared to infected young WT mice (Figure 4C). CD247 is an essential factor for the assembly of the TCR and the downstream signaling after stimulation with an antigen (Jin, Yuan, Mehta, Ezenwa, & Morel, 2024; Pitcher et al., 2003). In young WT mice gene expression levels of *CD247* were increased with infection, likely due to proliferation and infiltration of T cells. (Figure 4D). Notably, in infected Terc^ko/ko^ mice, the expression of *CD247* is absent. Moreover, there is no difference in *CD247* expression between non-infected and infected old WT mice (Figure 4D). Similar trends can be seen for other T cell markers, such as Programmed Cell Death 1 (*PDCD1*) and *CD4* (Figure 4E and F). Additionally, the TCR signaling pathway and Th1 and Th2 differentiation pathway are among the most enriched pathways in infected young WT mice compared to non-infected young WT mice. In contrast, those pathways are not upregulated when comparing infected Terc^ko/ko^ with non-infected Terc^ko/ko^ mice (Supplemental Figure 4A).

These findings indicate a malfunction in the activation of T cells as components of the TCR signaling pathway were partially downregulated. The complete absence of *CD247*, an essential factor for downstream activation of the TCR, indicates aberrant T cell function of Terc^ko/ko^ mice during infection.

### T cells of Terc^ko/ko^ mice do not adequately react to infectious challenge

To investigate the functionality of T cells and their CD247 expression in Terc^ko/ko^ mice during infection, murine T cells and AMs were isolated. An overactive inflammatory response could be a potential explanation for the dysregulated TCR signaling. Therefore, to study the impact of AMs on T cells we infected the co-culture of both isolated cells with a multiplicity of infection 5 (MOI5) of heat-killed (HK) *S. aureus* for 24 hours (Figure 5A). All components of the experimental setup were visualized using SEM. In particular, the close association between AMs and T cells was notable (Figure 5B). IF staining of AMs of Terc^ko/ko^ and young WT mice revealed the successful internalization of *S. aureus* as well as a change of phenotypic appearance to a more activated shape (Figure 5C).

**Figure 5:**
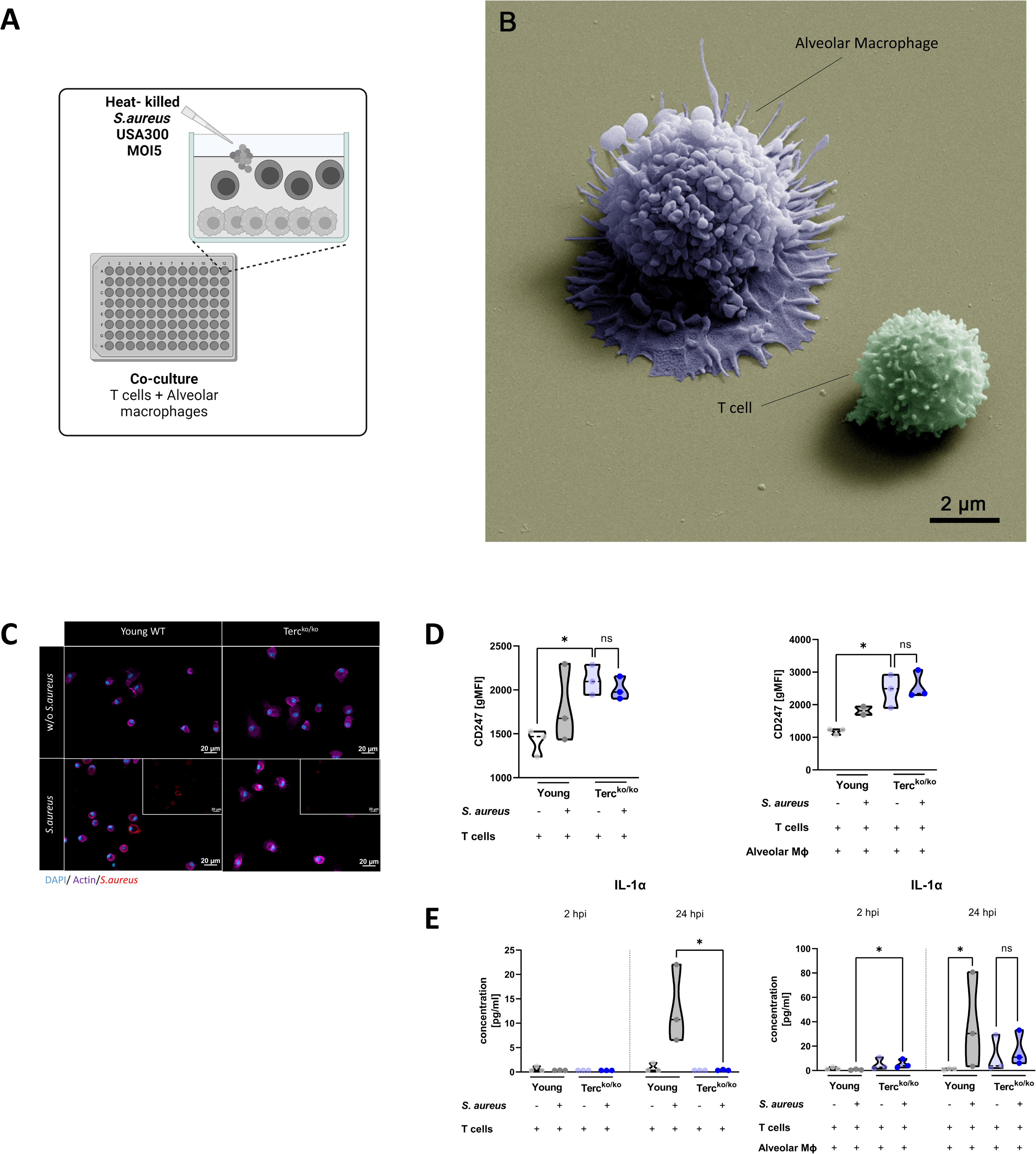
T cells of Terc^ko/ko^ mice do not adequately react to bacterial challenge. (A) Schematic representation of experiment. Murine T cells and AMs were isolated, co-cultured, and stimulated with heat-killed *S. aureus* with an MOI5 for 24 hours. (B) Colorized SEM picture of the experimental setup. An AM close to a T cell, is shown. (C) Immunofluorescence staining of infected and non-infected AMs from young WT and Terc^ko/ko^ mice. AMs were stained with antibodies for *S. aureus* (red), actin (purple), and DAPI (blue). (D) CD247 expression measured with flow cytometry of non-infected and infected T cells (left) and co-cultured T cells and AMs (right) of young WT (each n=3) and Terc^ko/ko^ mice (each n=3) after 24 hpi. CD247 expression is displayed as geometric mean fluorescence intensity (gMFI). (E) IL-1α from supernatants of T cells (left) and co-cultured T cells and AMs (right) of young WT (each n=3) and Terc^ko/ko^ mice (each n=3) at 2 and 24 hpi. IL-1α levels were measured using a flow cytometry based LEGENDPlex^TM^ mouse T helper cytokine panel. ns ≥ 0.05, *p < 0.05. P calculated by Kruskal-Wallis-test (D-E). Data is displayed as violin plot showing the median as well as lower and upper percentile of each dataset. Each replicate is displayed as a data point.

Utilizing flow cytometry, we observed a substantial elevation in the baseline expression of CD247 in T cells from Terc^ko/ko^ mice in comparison to T cells from young WT mice at 24 hpi (Figure 5D). However, there was no increase in CD247 expression upon stimulation with *S. aureus* in the Terc^ko/ko^ T cells (Figure 5D). In contrast, observation of T cells of young WT mice revealed a tendency towards an increase in CD247 expression in response to *S. aureus* stimulation (Figure 5D). This indicated improper activation of TCR signaling after *S. aureus* stimulation in T cells of Terc^ko/ko^ mice.

We could also detect differences in cytokine production when comparing the AMs and T cells from young WT and Terc^ko/ko^ mice. At 24 hpi, IL-1α production was significantly higher in T cells from young WT mice stimulated with *S. aureus* when compared to the T cells from Terc^ko/ko^ mice (Figure 5D, left). Notably, in the co-culture, IL-1α production was significantly higher in Terc^ko/ko^ cells 2 hpi (Figure 5D, right). On the contrary, there was a significant induction of IL-1α in the young WT co-culture but no increase in the Terc^ko/ko^ co-culture with *S. aureus* stimulation after 24 hours (Figure 5D, right).

Our data indicate that T cells from Terc^ko/ko^ mice do not effectively react to the infectious stimulus and thus seem dysfunctional due to TCR downregulation.

## Discussion

Shortening of telomeres is a key factor in the aging process and has a significant impact on other hallmarks of aging, such as mitochondrial dysfunction and cellular senescence (Birch, Barnes, & Passos, 2018; López-Otín et al., 2023). In this context, the discussion surrounding the importance of Terc is primarily confined to its role within telomerase. Consequently, other potential functions of Terc in various cellular processes have been largely overlooked. Interestingly, various studies have linked Terc to inflammation, as it has been shown to activate the NF-κB and PI3K-AKT pathways (Liu et al., 2019; Wu et al., 2022). Another study could show that the deletion of *T*erc in mice resulted in NLRP3 inflammasome activation after infection with *S. aureus*, thus contributing to more severe pneumonia (Kang et al., 2018). These studies suggest a close association between Terc, inflammation, and the response to bacterial lung infections.

In our study, we utilized a Terc^ko/ko^ *S. aureus* pneumonia model to investigate this connection. In accordance with German animal welfare regulations and the 3R principle, we only had a reduced number of Terc^ko/ko^ mice available for the experiment, as some infections were fatal. This reduced number of mice is a limitation of our study. However, we could demonstrate that Terc^ko/ko^ mice presented with a more severe pneumonia, higher bacterial load, and increased mortality compared to old and young WT mice (Figure 1). Additionally, we could show that infected Terc^ko/ko^ mice are rather heterogeneous and display several degrees of severity, which we investigated separately. Of 5 infected mice, 2 died, 2 displayed signs of a systemic infection and only 1 had a mild infection. The mice with systemic infection were grouped based on the presence of bacteria in organs other than the lung, as well as the sequencing data, where they clustered separately. The detection of bacteria in extra pulmonary organs is of particular interest, as bacteremia is a symptom of severe pneumonia and is associated with high mortality (De la Calle et al., 2016). The in this study conducted subclassification into different degrees of severity of Terc^koko^ mice is lacking in similar studies, highlighting the here gained detailed insights. Terc^ko/ko^ mice were previously shown to have higher susceptibility to *S. aureus* pneumonia, although they exhibited a reduced bacterial load in their lungs (Kang et al., 2018). These differences may be attributed to the missing separation of infected Terc^ko/ko^ mice into different degrees of severity. Additionally, this heterogeneity in disease manifestation points towards the limitations of mouse models to study *S. aureus* pathologies. There are a variety of differences between *S. aureus* infections in humans and mice. For instance several relevant virulence factors (e.g. Pantone-Valentine leukocidine) function less efficient in mice than in humans (Zheng et al., 2023). Therefore, higher infection doses are needed to successfully establish an infection that is comparable to humans (Zheng et al., 2023).

Inflammation and the activation of the NLRP3 inflammasome are necessary components to fight bacterial infections of the lung. However, excessive inflammation and over-activation of the NLRP3 inflammasome lead to lung injury and can impede pathogen clearance, thus contributing to *S. aureus*-induced pathologies (McVey, Steinberg, & Goldenberg, 2021; Wang et al., 2022). Upon examining the lungs of infected Terc^ko/ko^ mice with systemic infection, we identified a marked increase in inflammation parameters (Figure 2). Furthermore, we discovered that genes associated with the NLRP3 inflammasome pathway were elevated in the infected Terc^ko/ko^ mice but not in the corresponding non-infected control cohort. As published previously, this implies that *Terc* knockout on its own does not result in heightened inflammation and inflammasome activation but rather requires an external trigger (Kang et al., 2018). Inflammasome activation in Terc^ko/ko^ mice was connected to mitochondria dysfunction and an elevated reactive oxygen species (ROS) production. Additionally, regulation of the inflammasome seems to be dysfunctional as TNAIFP3, an important negative regulator of the inflammasome was downregulated in the Terc^ko/ko^ mice in this study (Kang et al., 2018). However, we did not observe downregulation of TNFAIP3, but rather an upregulation in infected Terc^ko/ko^ mice (Figure 2C). These differences may however be attributed to the different study design. Kang et al. focused on telomere dysfunction studying macrophages of old Terc^ko/ko^ mice, while our study investigated the entire lung of young Terc^ko/ko^ mice. Of particular note were the heightened levels of IL-1β, a factor known for its ability to amplify the inflammatory response and facilitate the infiltration of pro-inflammatory immune cells, thereby fostering pulmonary injury (Chen et al., 2018; Hu et al., 2020). Additionally, pro-inflammatory macrophages infiltrating the lung are critical to inflammation-mediated lung injury (Chen et al., 2018; Guan, Zhou, Zhou, & Lin, 2023). In our study, we could identify the upregulation of several pro-inflammatory macrophage markers in infected Terc^ko/ko^ mice, as well as several chemokine markers further exacerbating the inflammation in the lung via attraction of immune cells (Figure 3). Notably, we did not observe upregulation of inflammation parameters or activation of the inflammasome in the infected old WT mice. Although *CD14* showed a slight increase in non-infected old WT mice compared to non-infected young WT mice, there was no upregulation of macrophage markers in the infected cohorts of these groups. Innate immune cells of aged individuals especially macrophages are associated with chronic low grade inflammation contributing to injury during infection (Duong et al., 2021). However studies could also show that macrophages of aged mice had a reduced response to stimulation of pathogen recognition receptors such as Toll-like receptor 2 (TLR2). This results in an initially diminished and delayed response of macrophages to infectious challenge, which could be a possible explanation for the lack of inflammation at 24 hpi (Boe et al., 2017; Hinojosa, Boyd, & Orihuela, 2009).

An overactive inflammatory response not only causes tissue damage but also leads to T cell dysfunction (Guan et al., 2023). A possible mechanism is the downregulation of CD247, which is the CD3ζ chain of the TCR. CD247/CD3ζ is necessary for the assembly of the receptor at the cell surface. Furthermore due to its multiple immunoreceptor tyrosine-based activation (ITAM) motifs, it is a central factor in the signal transduction (Bronstein-Sitton et al., 2003; Jin et al., 2024). The downregulation or absence of CD247/CD3ζ is linked to reduced responsiveness in T cells and is commonly present in autoimmune diseases like rheumatoid arthritis and lupus (Dexiu, Xianying, Yingchun, & Jiafu, 2022). This mechanism also occurs in various chronic infectious diseases, including HIV and Hepatitis C (Dexiu et al., 2022). However, it can also be a general regulatory process in T cells during infection, helping to maintain immune homeostasis and prevent excessive inflammation (Baniyash, 2004; van der Donk et al., 2021).Thus, the connection between CD247/CD3ζ downregulation during chronic inflammation and chronic infectious diseases has been well documented (Dexiu et al., 2022). Interestingly, we could show that expression of *CD247* is entirely absent in infected Terc^ko/ko^ mice, while in the non-infected Terc^ko/ko^ mice expression of *CD247* could be detected after 24 hours of infection. Hence, our data indicates that infection combined with deletion of *Terc* is associated with reduction of *CD247* expression. Our study provides the first evidence of the total absence of *CD247* expression during acute bacterial infection and inflammation (Figure 4).

Previous work has already reported that constant stimulus with bacterial antigens reduces the expression of CD247/CD3ζ and impairs T cell function (Bronstein-Sitton et al., 2003). However, the complete absence of CD247/CD3ζ after a single antigenic stimulus, without chronic inflammation as well as the connection to knockout of *Terc*, are novel findings. These findings require further studies and investigations to gain a deeper understanding of the underlying mechanisms.

Downregulation of CD247/CD3ζ is facilitated by myeloid suppressor cells (MSC), which accumulate at the site of inflammation (Baniyash, 2004; Dexiu et al., 2022; Ezernitchi et al., 2006). MSC are cells which exert an immunosuppressive function on T cells in response to inflammation or infection (Medina & Hartl, 2018). Thus MSC contribute towards the regulation of the immune response to pathogens to prevent an overshooting reaction of the immune system (Medina & Hartl, 2018). However their immunosuppressive function can also exacerbate infections. For instance, elevated levels of MSC induced T cell dysfunctionality and caused an exacerbated and sustained chronic infection in an *S. aureus* mouse model (Tebartz et al., 2015). Immunosuppression of T cells is conveyed by multiple mechanisms, one of which is the downregulation of CD247/CD3ζ (Ezernitchi et al., 2006). This process is facilitated among others by the ability of MSCs to metabolize the amino acid arginine, as arginine seems to be essential for expression of CD247 in T cells (Ezernitchi et al., 2006; Rodriguez et al., 2004). Interestingly, we could identify several MSC markers, such as lymphocyte antigen 6 family member G (*LY6G)*, to be upregulated in the infected Terc^ko/ko^ mice. This suggests a potential role of MSCs in the downregulation of CD247/CD3ζ (Supplemental Figure 4D). Notably, we could identify reduced expression of *CD247* in infected old WT mice compared to young WT mice. Moreover, our sequencing data did not show an increase in *CD247* expression in infected old WT mice compared to the non-infected control group, which further suggests that the T cells of old WT mice may also exhibit some level of dysfunction (Figure 4). One potential underlying mechanism is the reduced ability of aged macrophages to activate T cells due to diminished expression of the receptor required for presentation of the antigen (Herrero, Marqués, Lloberas, & Celada, 2001). Furthermore, T cells of aged individuals become increasingly deficient due to immunosenescence and exhaustion (Mittelbrunn & Kroemer, 2021). Supporting this, we could observe downregulation of *CD27* compared to infected Young WT mice, while *CTLA4* was upregulated in infected old WT compared to uninfected mice (Supplemental Figure 2E and F). Downregulation of CD28 and CD27 is part of T cell aging and an essential marker of T cell senescence, while upregulation of CTLA4 is part of a signature pointing to T cell exhaustion (Zhao, Shao, & Peng, 2020). Interestingly, senescent T cells have shorter telomeres and exhibit reduced telomerase activity, which indicates a possible connection between Terc and T cell function (Mittelbrunn & Kroemer, 2021). Interestingly, *CD28* expression was downregulated in infected Terc^ko/ko^ mice compared to old WT infected mice (Figure 4A and C). However, we did not observe any other relevant changes of T cell senescence or exhaustion markers in infected Terc^ko/ko^ mice (Figure 4A and C). Thus T cell senescence could be a contributing factor to the T cell dysfunctionality, but likely does not explain the complete phenotype. The differences in the expression of T cell senescence and exhaustion markers, in addition to the lack of increased inflammation in old WT mice, indicates that mechanisms causing downregulation of parts of the TCR and T helper cell markers in old WT and Terc^ko/ko^ mice differ from each other. As T cell senescence is however a known consequence of *Terc* deletion, the lack of in depth investigation of this process in the present study is a limitation (Matthe et al., 2022). It is important to note that telomere shortening has a significant impact on the immune system (Chakravarti et al., 2021). Although young Terc^ko/ko^ mice were used in this study, telomere shortening is still likely to be a contributing factor. Therefore, further experiments investigating the role of T cell senescence in this model should be conducted.

Nevertheless, old WT mice exhibit certain immune cell dysfunction similarities. However, given the intricate nature of aging, other mechanisms contribute to the distinct response observed in Terc^ko/ko^ and old WT mice following infection. This highlights the necessity of employing a knockout model to specifically examine the influence of Terc on infection, isolating it from the multifaceted processes involved in aging.

We observed an increase in pro-inflammatory macrophages in the lungs of Terc^ko/ko^ mice during infection. To explore the role of macrophages in regulating T cell function during infection, we investigated whether AMs were sufficient to induce CD247/CD3ζ downregulation in T cells. While CD247/CD3ζ expression increased with bacterial stimulation in T cells from young WT mice, we failed to induce such an increase in CD247/CD3ζ expression in T cells from Terc^ko/ko^ mice (Figure 5). The expression pattern of CD247/CD3ζ was similar to that observed in the sequencing data, as initial levels of CD247/CD3ζ expression in non-infected Terc^ko/ko^ mice were higher than those of young WT mice. However, there was no significant reduction of CD247/CD3ζ after stimulation with bacteria in the T cells of Terc^ko/ko^ mice. One potential explanation for this finding is the absence of MSCs and other infiltrating cells, which can contribute to the inflammatory environment and lead to T cell dysfunction. Furthermore, after 24 but not 2 hours of infection, we observed a reduced release of the pro-inflammatory cytokine IL-1α from Terc^ko/ko^ T cells, suggesting a time-dependent reduction of T cell function. The results of the experiment conducted on T cells of Terc^ko/ko^ mice revealed an insufficient response to the bacterial stimulation, as there was no significant increase in CD247/CD3ζ expression or IL-1α release after 24 hpi. As total CD4^+^ T cells were analyzed in this study, it would be useful to investigate specific T cell populations such as memory and effector T cells to elucidate the potential mechanism leading to T cell dysfunctionality in further detail. Additionally, analysis of differences in immune cell recruitment to the lungs between young WT and Terc^ko/ko^ mice would be relevant. Thus, additional experiments are needed to validate and further investigate the underlying mechanisms leading to T cell dysfunctionality in Terc^ko/ko^ mice.

Our data points towards the pivotal role of Terc in the regulation of inflammation as well as maintaining immune homeostasis during infection. We could demonstrate that *S. aureus* pneumonia in Terc^ko/ko^ mice resulted in an overshooting inflammation and activation of the NLRP3 inflammasome. The elevated inflammation eventually leads to T cell dysfunction, which impairs the host’s ability to mount an effective immune response against the pathogen, resulting in increased bacterial load and mortality. As T cell dysfunctionality is caused by the excessive inflammation, other pathogens inducing high inflammatory responses in the host could also lead to a similar phenotype in Terc^ko/ko^ mice.

This study linked T cell dysfunction to Terc deletion for the first time and thus provided insights into possible mechanisms contributing to a more severe *S. aureus* pneumonia in the aging population.

## Acknowledgments

Breeding pairs of Terc^ko/ko^ mice were kindly provided by Lenhard Rudolph (Leibniz Institute on Aging, Jena, Germany). We thank Sylvia Hänßgen, Yvonne Ozegowski, and Lea Herrmann for their excellent technical assistance.

## Author Contributions

Conceptualization, S.D.-E.; methodology, Y.R; F.H., A.H., T.L., S.N., writing—original draft preparation, Y.R.; S.D.-E. writing—review and editing, all authors; visualization: Y.R., S.D.-E.; resources and supervision: B.L., S.D.-E.; funding acquisition, B.L.; S.D.-E. All authors have read and agreed to the published version of the manuscript.

## Data availability statement

The data supporting this study’s findings are available from the corresponding author upon reasonable request.

---

**Supplemental Figure 1: Terc^ko/ko^ mice with systemic infection display more severe pneumonia with bacteremia and increased mortality.**

(A) Clinical score of infected Terc^ko/ko^ mice at the time of infection (0 dpi) and 24 hpi (1 dpi). The mice were divided into three groups according to their clinical score values: mice with mild infection (dark blue dots; n=1), mice with systemic infection (light blue dots; n=2), and mice where infection resulted in a fatal outcome (black dots; n=2). Mice with systemic infection were defined as mice with an increased clinical score and the presence of bacteria in organs other the lung.

(B) Bacterial load in kidney and liver tissue of the infected Terc^ko/ko^ mice 24 hpi (each n=3). Light blue data points represent mice with systemic infection, while dark blue dots represent the mouse with mild infection.

(C) Bacterial load of the lung of infected mice at 24 hpi. Young WT and old WT mice: n=5; Terc^ko/ko^ mice with systemic infection: n=2, Terc^ko/ko^ mice without systemic infection: n=1. Data is displayed as logarithmic.

(D) Relative body weight of young WT (each n=5), old WT (each n=5) and Terc^ko/ko^ mice (non-infected: n = 3; with systemic infection: n=2; without systemic infection: n =1). Relative body weight displays body weight at 24 hpi as a percentage of the body weight at the time of infection.

(E) Relative lung weight of y young WT (each n=5), old WT (each n=5) and Terc^ko/ko^ mice (non-infected: n = 3; with systemic infection: n=2; without systemic infection: n =1). Relative lung weight displays lung weight after 24 hpi as a percentage of the current body weight.

(F) Measurement of average telomere length per chromosome end in lungs of young WT (n=4), old WT (n=4) and Terc^ko/ko^ mice (n=4).

(G) PCA of sequenced lungs from non-infected and infected youngWT (each n=3), old WT (each n=3) and Terc^ko/ko^ mice (each n=3). The mice with systemic infection clustered separately from the other samples. Nevertheless, the infected Terc^ko/ko^ mice were considered one group for the expression analysis and not split into separate groups for the subsequent analysis.

(H) Number of DEGs per comparison including up- and downregulated DEGs as well as the total count. Differential gene expression analysis was conducted using the DESeq2 software as well as negative binomial distribution. For FDR calculation the Benjamini-Hochberg procedure was used. Sequencing data displays mRNA sequencing data of non-infected and infected lungs of the three different mice cohorts at 24 hpi. For each mouse cohort three biological replicates were sequenced and used for analysis.

Infected Terc^ko/ko^ mice were divided into systemic and non-systemic infected mice for several analyses (C, D, and E). Data is displayed as violin plot showing the median as well as lower and upper percentile of each dataset. *p < 0.05. P calculated by Kruskal-Wallis-test (C-E). Each replicate is displayed as a data point.

**Supplemental Figure 2: Several pathways relevant for inflammation and adaptive and innate immune response are differentially expressed in infected Terc^ko/ko^ mice.**

Heatmaps of DEGs related to the NLRP3 inflammasome pathway (A), macrophage marker (C), Chemokine signaling pathway (D), T helper cell marker (E), or TCR signaling (F). Red indicates upregulation, and blue indicates downregulation of the gene. Differential gene expression analysis was conducted using the DESeq2 software as well as negative binomial distribution. For FDR calculation the Benjamini-Hochberg procedure was used. Heatmaps were constructed using R Studio. Only significant DEGs (p-value < 0.05) were displayed in the heatmap. Non-significant genes were set to zero and are shown as white. DEGs are displayed as log2fold-change. For each mouse cohort three biological replicates were sequenced and used for analysis. The respective groups used for the comparison are indicated below each column. All comparisons are shown. All heatmaps display mRNA sequencing data of non-infected and infected lungs of the three different mice cohorts at 24 hpi.

B) Immunofluorescence staining of CD68 (red), actin (purple), and DAPI (blue) of AMs isolated from Terc^ko/ko^ and young WT mice. Representative pictures are shown for each group.

**Supplemental Figure 3:**

(A) Exemplary gating strategy for the immunophenotyping of cells from murine spleen. First, lymphocytes were selected, then duplicates and dead cells were excluded (APC-Cy7 negative). To avoid autofluorescence of macrophages, the F4/80 negative population was selected. For further calculations of T cell frequencies the CD3^+^ CD19^-^ cell population was used. Other immune cell populations were selected by the indicated marker. For calculation of macrophage frequencies the F4/80 positive population was selected.

B) Percentage of Ly-6G^+^ -living cells in the spleen of non-infected young WT (n=3), old WT (n=3), and Terc^ko/ko^ mice (n=5). Cells were analyzed with flow cytometry.

C) Percentage of CD19^+^ -living cells in the spleen of non-infected young WT (n=3), old WT (n=3), and Terc^ko/ko^ mice (n=5). Cells were analyzed with flow cytometry.

D) Percentage of CD335^+^ -living cells in the spleen of non-infected young WT (n=3), old WT (n=3), and Terc^ko/ko^ mice (n=5). Cells were analyzed with flow cytometry.

E) Percentage of F4/80^+^ -living cells in the spleen of non-infected young WT (n=3), old WT (n=3), and Terc^ko/ko^ mice (n=5). Cells were analyzed with flow cytometry.

F) Percentage of CD11c^+^ -living cells in the spleen of non-infected young WT (n=3), old WT (n=3), and Terc^ko/ko^ mice (n=5). Cells were analyzed with flow cytometry.

ns ≥ 0.05, *p < 0.05, **p < 0.01. P calculated by Kruskal-Wallis-test (B-F). Data is displayed as violin plot showing the median as well as lower and upper percentile of each dataset. Each replicate is displayed as a data point.

**Supplemental Figure 4:**

(A) KEGG pathway enrichment analysis comparing infected Terc^ko/ko^ vs. infected old WT, infected young WT vs. non-infected young WT and infected Terc^ko/ko^ vs. non-infected Terc^ko/ko^ mice. The gene ratio describes the ratio of differentially expressed genes to all genes of the respective pathway. The dot plots were constructed using R Studio.

(B) Gating strategy for T cells isolated from murine spleen to asses CD247 expression. First, lymphocytes were selected, duplicates and dead cells were excluded before selecting CD3^+^ CD247^+^ cells. This population was used for further calculations.

(C) CD44 expression measured with flow cytometry of non-infected and infected T cells (left) and co-cultured T cells and AMs (right) of young WT (each n=3) and Terc^ko/ko^ mice (each n=3) after 24 hpi. CD44 expression is displayed as geometric mean fluorescence intensity (gMFI).

(D) Heatmap of differentially regulated MSC markers. Red indicates upregulation, and blue indicates downregulation of the gene. Differential gene expression analysis was conducted using the DESeq2 software as well as negative binomial distribution. For FDR calculation the Benjamini-Hochberg procedure was used. Heatmaps were constructed using R Studio. Only significant DEGs (p-value < 0.05) were displayed in the heatmap. Non-significant genes were set to zero and are shown as white. DEGs are displayed as log2fold-change. For each mouse cohort three biological replicates were sequenced and used for analysis. The respective groups used for the comparison are indicated below each column. All heatmaps displays mRNA sequencing data of non-infected and infected lungs of the three different mice cohorts at 24 hpi.

ns ≥ 0.05. P calculated by Kruskal-Wallis-test (C). Data is displayed as violin plot showing the median as well as lower and upper percentile of each dataset. Each replicate is displayed as a data point.

**Supplemental Figure 5:**

(A) Overview of the methods used in this publication. Illustration was created with Biorender.

**Supplemental Figure 6:**

(A) Exemplary results of genotyping of Terc^ko/ko^ mice used for breeding. Knockout mice show a single band at 180 bp, while WT mice have a single band at 200 bp. Heterozygous mice (WT/ko) display both bands. Results were analyzed using an Agilent 2200 TapeStation.

## References

Baniyash, M. (2004). TCR ζ-chain downregulation: curtailing an excessive inflammatory immune response. Nature Reviews Immunology, 4(9), 675–687. doi:10.1038/nri1434

Birch, J., Barnes, P. J., & Passos, J. F. (2018). Mitochondria, telomeres and cell senescence: Implications for lung ageing and disease. Pharmacology & Therapeutics, 183, 34–49. 10.1016/j.pharmthera.2017.10.005

Boe, D. M., Boule, L. A., & Kovacs, E. J. (2017). Innate immune responses in the ageing lung. Clin Exp Immunol, 187(1), 16–25. doi:10.1111/cei.12881

Bronstein-Sitton, N., Cohen-Daniel, L., Vaknin, I., Ezernitchi, A. V., Leshem, B., Halabi, A., … Baniyash, M. (2003). Sustained exposure to bacterial antigen induces interferon-γ-dependent T cell receptor ζ down-regulation and impaired T cell function. Nature Immunology, 4(10), 957–964. doi:10.1038/ni975

Canan, C. H., Gokhale, N. S., Carruthers, B., Lafuse, W. P., Schlesinger, L. S., Torrelles, J. B., & Turner, J. (2014). Characterization of lung inflammation and its impact on macrophage function in aging. Journal of Leukocyte Biology, 96(3), 473–480. doi:10.1189/jlb.4A0214-093RR

Chakravarti, D., LaBella, K. A., & DePinho, R. A. (2021). Telomeres: history, health, and hallmarks of aging. Cell, 184(2), 306–322. 10.1016/j.cell.2020.12.028

Chen, J., Wang, S., Fu, R., Zhou, M., Zhang, T., Pan, W., … Huang, Y. (2018). RIP3 dependent NLRP3 inflammasome activation is implicated in acute lung injury in mice. Journal of Translational Medicine, 16(1), 233. doi:10.1186/s12967-018-1606-4

De la Calle, C., Morata, L., Cobos-Trigueros, N., Martinez, J. A., Cardozo, C., Mensa, J., & Soriano, A. (2016). Staphylococcus aureus bacteremic pneumonia. European Journal of Clinical Microbiology & Infectious Diseases, 35(3), 497–502. doi:10.1007/s10096-015-2566-8

Dexiu, C., Xianying, L., Yingchun, H., & Jiafu, L. (2022). Advances in CD247. Scand J Immunol, 96(1), e13170. doi:10.1111/sji.13170

Duong, L., Radley, H. G., Lee, B., Dye, D. E., Pixley, F. J., Grounds, M. D., … Jackaman, C. (2021). Macrophage function in the elderly and impact on injury repair and cancer. Immunity & Ageing, 18(1), 4. doi:10.1186/s12979-021-00215-2

Ezernitchi, A. a. V., Vaknin, I., Cohen-Daniel, L., Levy, O., Manaster, E., Halabi, A., … Baniyash, M. (2006). TCR ζ Down-Regulation under Chronic Inflammation Is Mediated by Myeloid Suppressor Cells Differentially Distributed between Various Lymphatic Organs1. The Journal of Immunology, 177(7), 4763–4772. doi:10.4049/jimmunol.177.7.4763

Franceschi, C., Bonafè, M., Valensin, S., Olivieri, F., De Luca, M., Ottaviani, E., & De Benedictis, G. (2000). Inflamm-aging. An evolutionary perspective on immunosenescence. Ann N Y Acad Sci, 908, 244–254. doi:10.1111/j.1749-6632.2000.tb06651.x

Greider, C. W., & Blackburn, E. H. (1985). Identification of a specific telomere terminal transferase activity in Tetrahymena extracts. Cell, 43(2 Pt 1), 405–413. doi:10.1016/0092-8674(85)90170-9

Guan, T., Zhou, X., Zhou, W., & Lin, H. (2023). Regulatory T cell and macrophage crosstalk in acute lung injury: future perspectives. Cell Death Discovery, 9(1), 9. doi:10.1038/s41420-023-01310-7

Herrero, C., Marqués, L., Lloberas, J., & Celada, A. (2001). IFN-gamma-dependent transcription of MHC class II IA is impaired in macrophages from aged mice. J Clin Invest, 107(4), 485–493. doi:10.1172/jci11696

Hinojosa, E., Boyd, A. R., & Orihuela, C. J. (2009). Age-associated inflammation and toll-like receptor dysfunction prime the lungs for pneumococcal pneumonia. J Infect Dis, 200(4), 546–554. doi:10.1086/600870

Hodes, R. J., Hathcock, K. S., & Weng, N.-p. (2002). Telomeres in T and B cells. Nature Reviews Immunology, 2(9), 699–706. doi:10.1038/nri890

Hohensinner, P. J., Goronzy, J. J., & Weyand, C. M. (2011). Telomere dysfunction, autoimmunity and aging. Aging Dis, 2(6), 524–537. Retrieved from https://www.ncbi.nlm.nih.gov/pmc/articles/PMC3295061/pdf/ad-2-6-524.pdf

Hornung, F., Schulz, L., Köse-Vogel, N., Häder, A., Grießhammer, J., Wittschieber, D., … Deinhardt-Emmer, S. (2023). Thoracic adipose tissue contributes to severe virus infection of the lung. International Journal of Obesity, 47(11), 1088–1099. doi:10.1038/s41366-023-01362-w

Howden, B. P., Giulieri, S. G., Wong Fok Lung, T., Baines, S. L., Sharkey, L. K., Lee, J. Y. H., … Stinear, T. P. (2023). Staphylococcus aureus host interactions and adaptation. Nature Reviews Microbiology, 21(6), 380–395. doi:10.1038/s41579-023-00852-y

Hu, X., Qin, H., Li, Y., Li, J., Fu, L., Li, M., … Yin, B. (2020). Biochanin A protect against lipopolysaccharide-induced acute lung injury in mice by regulating TLR4/NF-κB and PPAR-γ pathway. Microbial Pathogenesis, 138, 103846. 10.1016/j.micpath.2019.103846

Jin, Y., Yuan, H., Mehta, I., Ezenwa, O., & Morel, P. A. (2024). Alternatively Spliced Variants of Murine CD247 Influence T Cell Development and Activation, Revealing the Importance of the CD3ζ C-Terminal Region. J Immunol, 212(4), 541–550. doi:10.4049/jimmunol.2300511

Jong, N. W. M. d., Kessel, K. P. M. v., & Strijp, J. A. G. v. (2019). Immune Evasion by Staphylococcus aureus. Microbiology Spectrum, 7(2), 10.1128/microbiolspec.gpp1123-0061-2019. doi:doi:10.1128/microbiolspec.gpp3-0061-2019

Kang, Y., Zhang, H., Zhao, Y., Wang, Y., Wang, W., He, Y., … Shi, L. (2018). Telomere Dysfunction Disturbs Macrophage Mitochondrial Metabolism and the NLRP3 Inflammasome through the PGC-1α/TNFAIP3 Axis. Cell Reports, 22(13), 3493–3506. 10.1016/j.celrep.2018.02.071

Lee, L.-N., Chou, W.-R., Wang, J.-Y., Kuo, Y.-L., Chang, C.-Y., Lee, Y.-C., … Liu, W.-L. (2022). Characteristics and local risk factors of community-acquired and health-care-associated Staphylococcus aureus pneumonia. Scientific Reports, 12(1), 18670. doi:10.1038/s41598-022-23246-1

Liu, H., Yang, Y., Ge, Y., Liu, J., & Zhao, Y. (2019). TERC promotes cellular inflammatory response independent of telomerase. Nucleic Acids Research, 47(15), 8084–8095. doi:10.1093/nar/gkz584

López-Otín, C., Blasco, M. A., Partridge, L., Serrano, M., & Kroemer, G. (2023). Hallmarks of aging: An expanding universe. Cell, 186(2), 243–278. 10.1016/j.cell.2022.11.001

Malainou, C., Abdin, S. M., Lachmann, N., Matt, U., & Herold, S. (2023). Alveolar macrophages in tissue homeostasis, inflammation, and infection: evolving concepts of therapeutic targeting. The Journal of Clinical Investigation, 133(19). doi:10.1172/JCI170501

Matthe, D. M., Thoma, O. M., Sperka, T., Neurath, M. F., & Waldner, M. J. (2022). Telomerase deficiency reflects age-associated changes in CD4+ T cells. Immun Ageing, 19(1), 16. doi:10.1186/s12979-022-00273-0

McVey, M. J., Steinberg, B. E., & Goldenberg, N. M. (2021). Inflammasome activation in acute lung injury. American Journal of Physiology-Lung Cellular and Molecular Physiology, 320(2), L165–L178. doi:10.1152/ajplung.00303.2020

Medina, E., & Hartl, D. (2018). Myeloid-Derived Suppressor Cells in Infection: A General Overview. J Innate Immun, 10(5-6), 407–413. doi:10.1159/000489830

Mittelbrunn, M., & Kroemer, G. (2021). Hallmarks of T cell aging. Nature Immunology, 22(6), 687–698. doi:10.1038/s41590-021-00927-z

Mogilenko, D. A., Shchukina, I., & Artyomov, M. N. (2022). Immune ageing at single-cell resolution. Nature Reviews Immunology, 22(8), 484–498. doi:10.1038/s41577-021-00646-4

Montecino-Rodriguez, E., Berent-Maoz, B., & Dorshkind, K. (2013). Causes, consequences, and reversal of immune system aging. The Journal of Clinical Investigation, 123(3), 958–965. doi:10.1172/JCI64096

Pitcher, L. A., Young, J. A., Mathis, M. A., Wrage, P. C., Bartók, B., & Van Oers, N. S. C. (2003). The formation and functions of the 21- and 23-kDa tyrosine-phosphorylated TCR ζ subunits. Immunological Reviews, 191(1), 47–61. 10.1034/j.1600-065X.2003.00003.x

Rios, F. J., Touyz, R. M., & Montezano, A. C. (2017). Isolation and Differentiation of Murine Macrophages. In R. M. Touyz & E. L. Schiffrin (Eds.), Hypertension: Methods and Protocols (pp. 297–309). New York, NY: Springer New York.

Rodriguez, P. C., Quiceno, D. G., Zabaleta, J., Ortiz, B., Zea, A. H., Piazuelo, M. B., … Ochoa, A. C. (2004). Arginase I Production in the Tumor Microenvironment by Mature Myeloid Cells Inhibits T-Cell Receptor Expression and Antigen-Specific T-Cell Responses. Cancer Research, 64(16), 5839–5849. doi:10.1158/0008-5472.Can-04-0465

Sahin, E., & DePinho, R. A. (2010). Linking functional decline of telomeres, mitochondria and stem cells during ageing. Nature, 464(7288), 520–528. doi:10.1038/nature08982

Shay, J. W., & Wright, W. E. (2019). Telomeres and telomerase: three decades of progress. Nature Reviews Genetics, 20(5), 299–309. doi:10.1038/s41576-019-0099-1

Tarry-Adkins, J. L., Aiken, C. E., Dearden, L., Fernandez-Twinn, D. S., & Ozanne, S. (2021). Exploring Telomere Dynamics in Aging Male Rat Tissues: Can Tissue-Specific Differences Contribute to Age-Associated Pathologies? Gerontology, 67(2), 233–242. doi:10.1159/000511608

Tebartz, C., Horst, S. A., Sparwasser, T., Huehn, J., Beineke, A., Peters, G., & Medina, E. (2015). A major role for myeloid-derived suppressor cells and a minor role for regulatory T cells in immunosuppression during Staphylococcus aureus infection. J Immunol, 194(3), 1100–1111. doi:10.4049/jimmunol.1400196

Torres, A., Cilloniz, C., Niederman, M. S., Menéndez, R., Chalmers, J. D., Wunderink, R. G., & van der Poll, T. (2021). Pneumonia. Nature Reviews Disease Primers, 7(1), 25. doi:10.1038/s41572-021-00259-0

van der Donk, L. E. H., Ates, L. S., van der Spek, J., Tukker, L. M., Geijtenbeek, T. B. H., & van Heijst, J. W. J. (2021). Separate signaling events control TCR downregulation and T cell activation in primary human T cells. Immun Inflamm Dis, 9(1), 223–238. doi:10.1002/iid3.383

Wang, Y., Cao, C., Zhu, Y., Fan, H., Liu, Q., Liu, Y., … Wu, M. (2022). TREM2/β-catenin attenuates NLRP3 inflammasome-mediated macrophage pyroptosis to promote bacterial clearance of pyogenic bacteria. Cell Death & Disease, 13(9), 771. doi:10.1038/s41419-022-05193-x

Wong, L. S. M., Oeseburg, H., de Boer, R. A., van Gilst, W. H., van Veldhuisen, D. J., & van der Harst, P. (2008). Telomere biology in cardiovascular disease: the TERC−/− mouse as a model for heart failure and ageing. Cardiovascular Research, 81(2), 244–252. doi:10.1093/cvr/cvn337

Wu, S., Ge, Y., Lin, K., Liu, Q., Zhou, H., Hu, Q., … Ju, Z. (2022). Telomerase RNA TERC and the PI3K-AKT pathway form a positive feedback loop to regulate cell proliferation independent of telomerase activity. Nucleic Acids Res, 50(7), 3764–3776. doi:10.1093/nar/gkac179

Zhang, M. W., Zhao, P., Yung, W. H., Sheng, Y., Ke, Y., & Qian, Z. M. (2018). Tissue iron is negatively correlated with TERC or TERT mRNA expression: A heterochronic parabiosis study in mice. Aging (Albany NY), 10(12), 3834–3850. doi:10.18632/aging.101676

Zhao, Y., Shao, Q., & Peng, G. (2020). Exhaustion and senescence: two crucial dysfunctional states of T cells in the tumor microenvironment. Cellular & Molecular Immunology, 17(1), 27–35. doi:10.1038/s41423-019-0344-8

Zheng, P., Liu, F., Long, J., Jin, Y., Chen, S., Duan, G., & Yang, H. (2023). Latest Advances in the Application of Humanized Mouse Model for Staphylococcus aureus. The Journal of Infectious Diseases, 228(6), 800–809. doi:10.1093/infdis/jiad253

