## Supplemental Figure 1-6 for "Telomerase RNA component knockout exacerbates *S. aureus* pneumonia by extensive inflammation and dysfunction of T cells"

SUPPLEMENTAL FIGURES

### Supplemental Figure 1

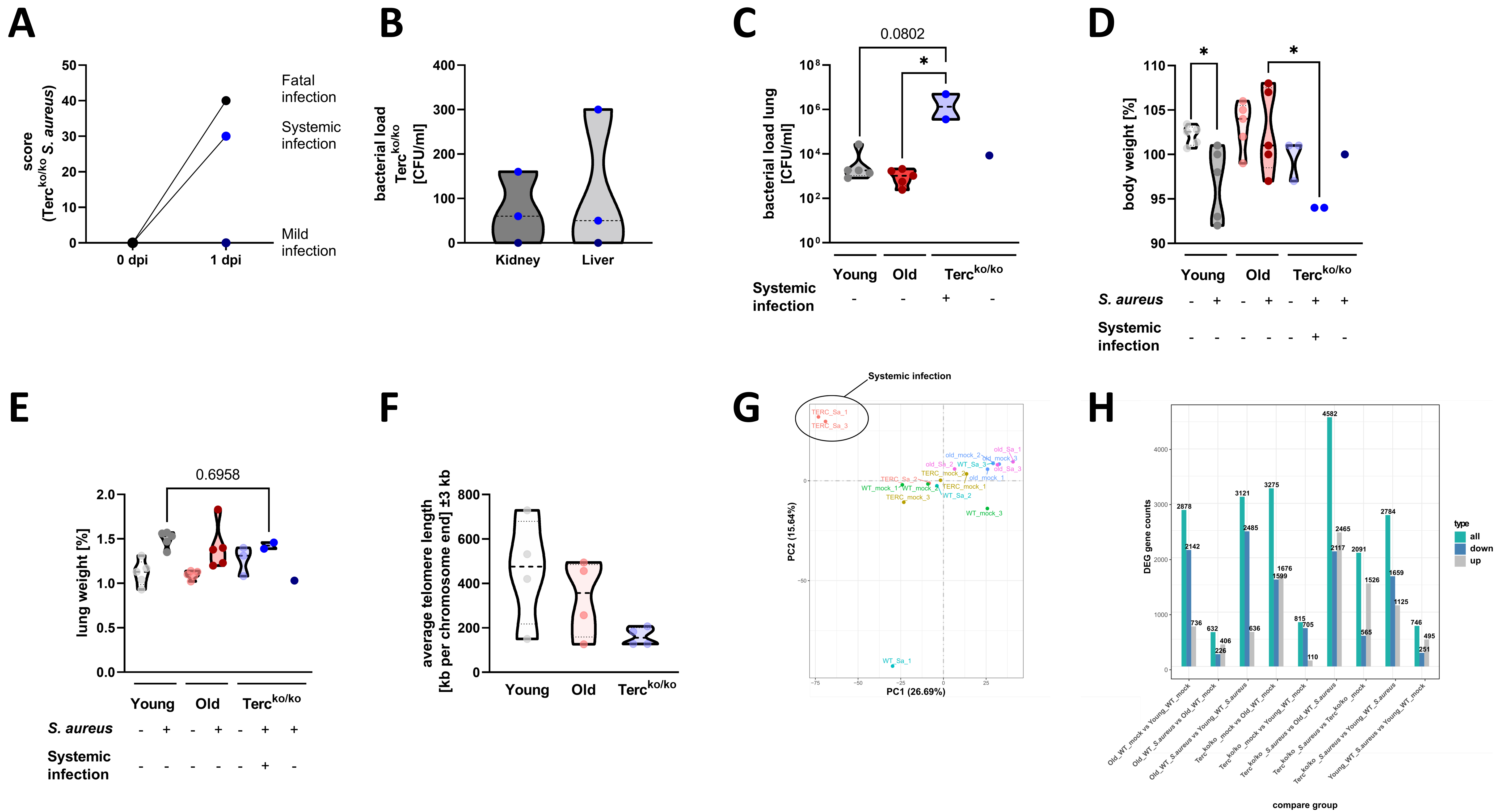

### Supplemental Figure 2

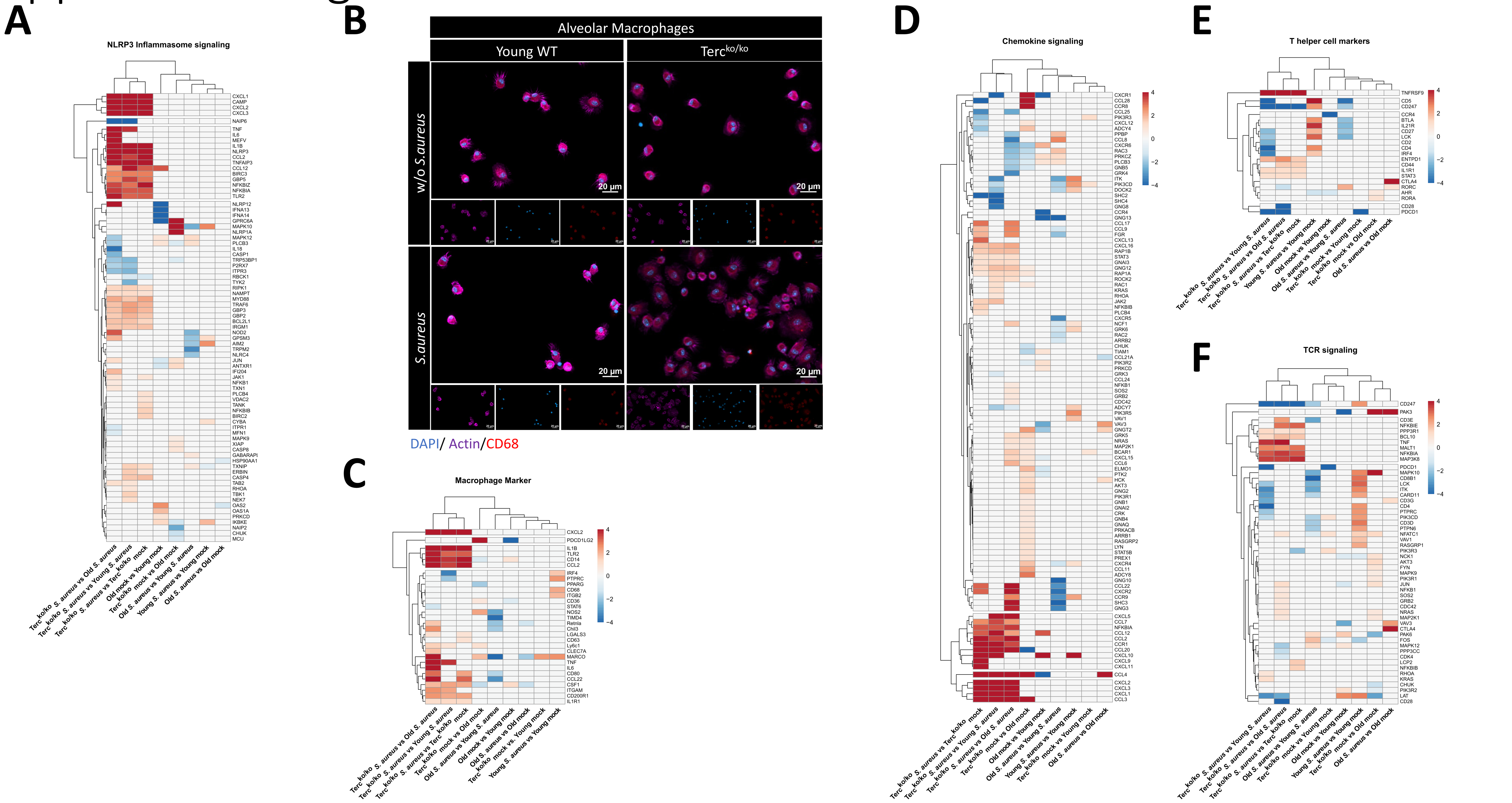

### Supplemental Figure 3

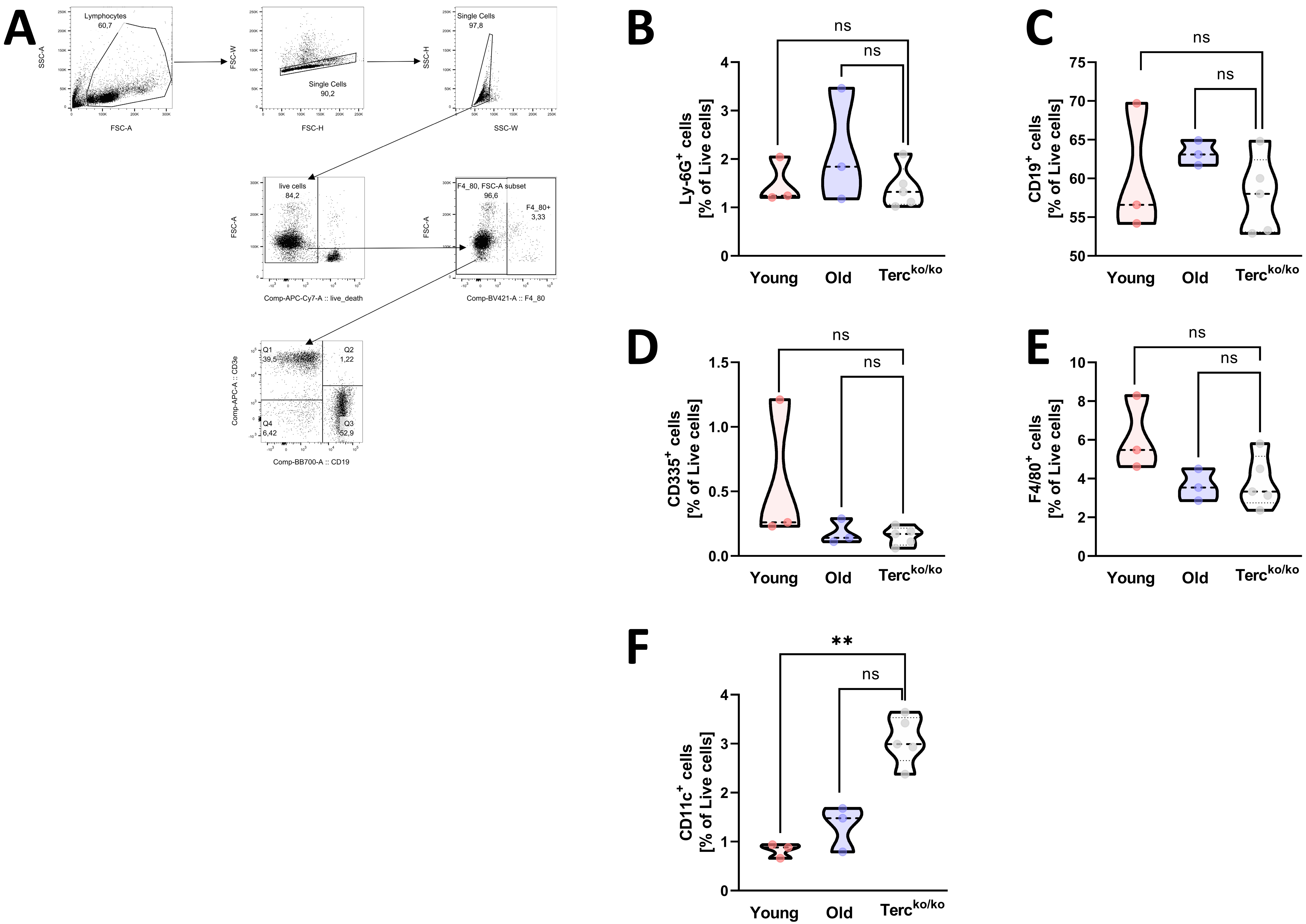

### Supplemental Figure 4

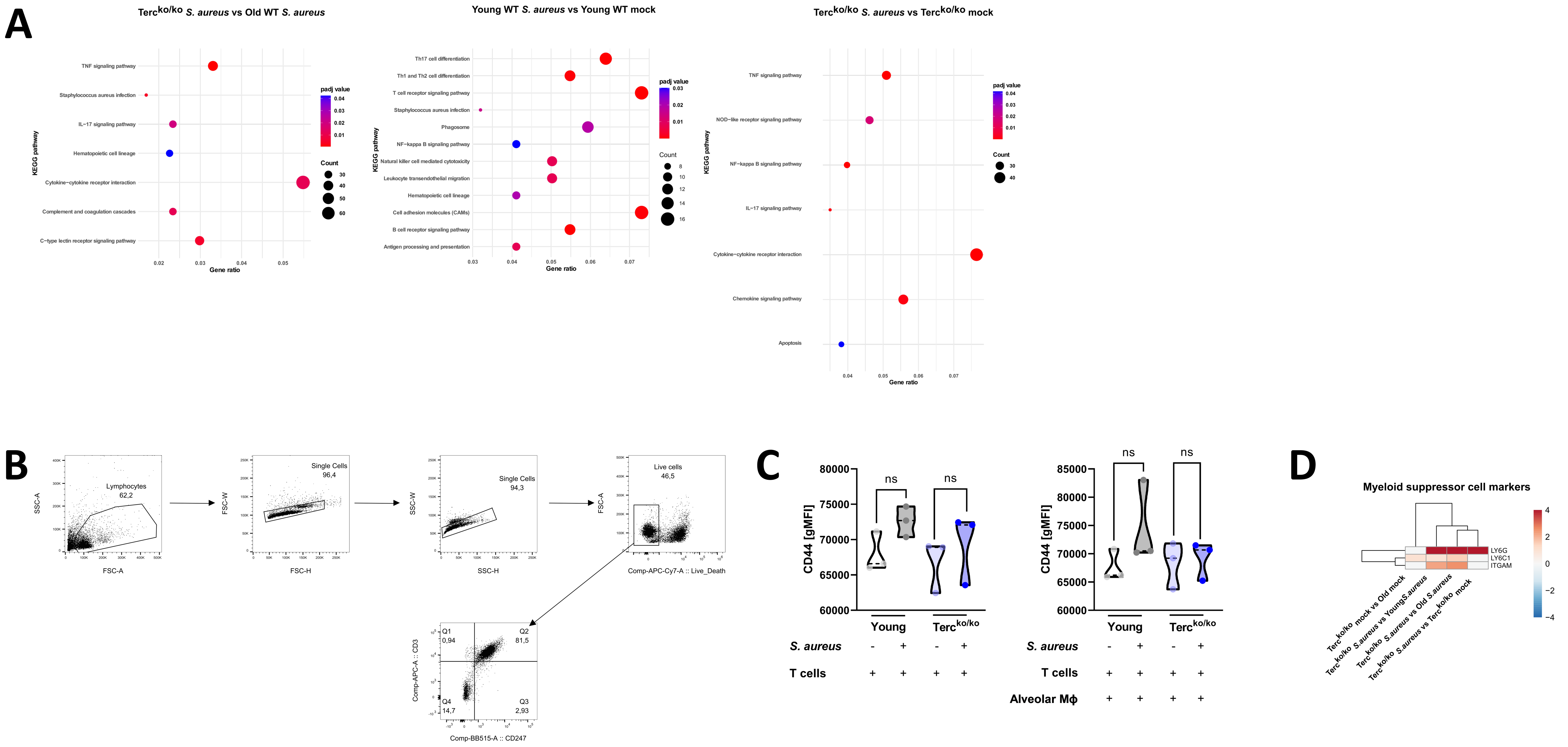

### Supplemental Figure 5

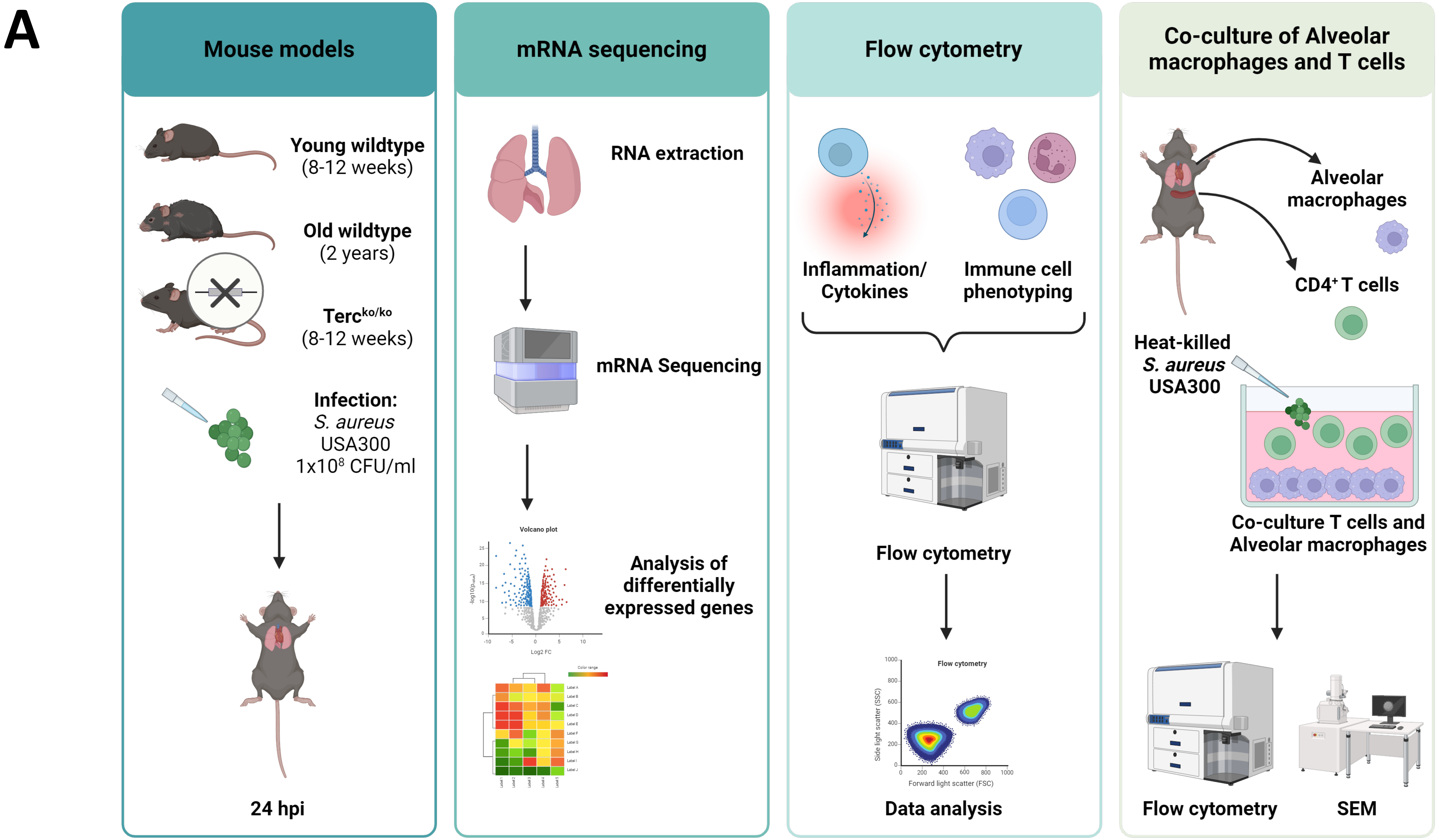

### Supplemental Figure 6

## A

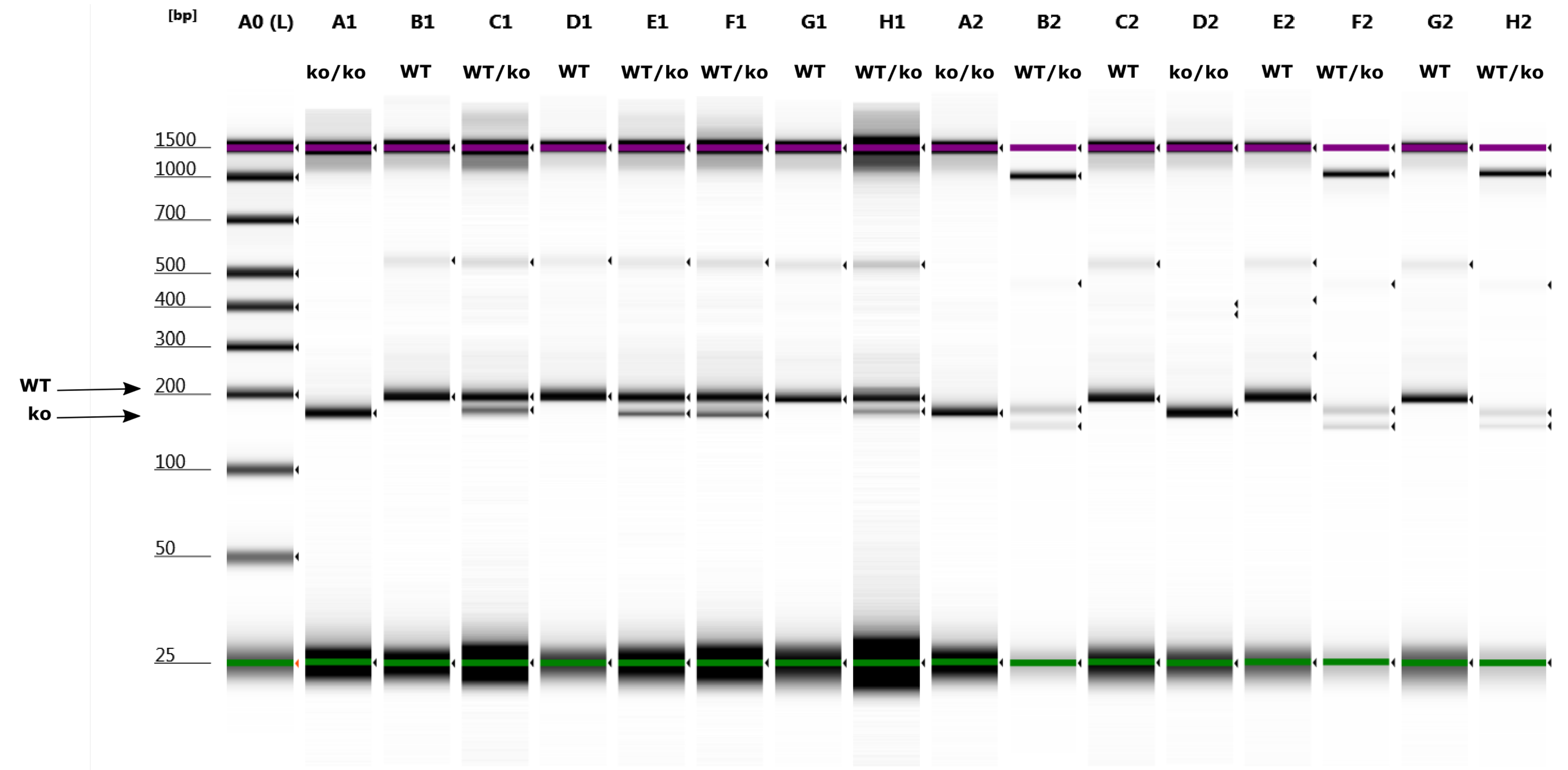


Default image (Contrast 69%), Image is Scaled to Sample

Sample Info

| Well | Conc. [ng/µl] | Sample Description | Alert | Observations |
|---|---|---|---|---|
| A0 | 20.3 | Electronic Ladder | | Ladder |
| A1 | 1.30 | 01 | | |
| B1 | 1.99 | 02 | | |
| C1 | 1.24 | 03 | | |
| D1 | 3.33 | 04 | | |
| E1 | 2.57 | 05 | | |
| F1 | 1.92 | 06 | | |
| G1 | 1.92 | 07 | | |
| H1 | 0.807 | 08 | | |
| A2 | 1.77 | 09 | | |
| B2 | 180 | 10 | | |
| C2 | 2.45 | 11 | | |
| D2 | 3.08 | 12 | | |
| E2 | 4.50 | 13 | | |
| F2 | 241 | 14 | | |
| G2 | 3.30 | 15 | | |
| H2 | 331 | 16 | | |
